# Direct binding of Cdt2 to PCNA is important for targeting the CRL4^Cdt2^ E3 ligase activity to Cdt1

**DOI:** 10.1101/468413

**Authors:** Akiyo Hayashi, Nickolaos Nikiforos Giakoumakis, Tatjana Heidebrecht, Takashi Ishii, Andreas Panagopoulos, Christophe Caillat, Michiyo Takahara, Richard G. Hibbert, Naohiro Suenaga, Magda Stadnik-Spiewak, Tatsuro Takahashi, Yasushi Shiomi, Stavros Taraviras, Eleonore von Castelmur, Zoi Lygerou, Anastassis Perrakis, Hideo Nishitani

## Abstract

The CRL4^Cdt2^ ubiquitin ligase complex is an essential regulator of cell-cycle progression and genome stability, ubiquitinating substrates such as p21, Set8 and Cdt1, via a display of substrate degrons on PCNA. Here, we examine the hierarchy of the ligase and substrate recruitment kinetics onto PCNA at sites of DNA replication. We demonstrate that the C-terminal end of Cdt2 bears a PCNA interaction protein motif (PIP box, Cdt2^PIP^), which is necessary and sufficient for binding of Cdt2 to PCNA. Cdt2^PIP^ binds PCNA directly with high affinity, two orders of magnitude tighter than the PIP box of Cdt1. X-ray crystallographic structures of PCNA bound to Cdt2^PIP^ and Cdt1^PIP^ show that the peptides occupy all three binding sites of the trimeric PCNA ring. Mutating Cdt2^PIP^ weakens the interaction with PCNA, rendering CRL4^Cdt2^ less effective in Cdt1 ubiquitination and leading to defects in Cdt1 degradation. The molecular mechanism we present suggests a new paradigm for bringing substrates to the CRL4-type ligase, where the substrate receptor and substrates bind to a common multivalent docking platform to enable subsequent ubiquitination.

**Summary blurb:** The C-terminal end of Cdt2 contains a PIP-box for binding to PCNA to promote CRL4^Cdt2^ function, creating a new paradigm, where the substrate receptor and substrates bind to a common multivalent docking platform for ubiquitination.

## INTRODUCTION

The integrity of genomic information is critical for proper cell function and cell survival, and is maintained by faithful replication during the S phase and segregation of duplicated chromosomes during mitosis. Cells are continuously challenged by genotoxic agents and environmental stress, and have complex mechanisms to activate DNA damage checkpoints, prevent cell-cycle progression, and repair the damaged DNA (Branzei and Foiani, 2010; Hoeijmakers, 2001). Many of the cell cycle transition events, as well as responses to DNA damage, are driven by E3 Cullin-RING ubiquitin Ligases (CRL), that catalyse the ubiquitination and destruction of specific protein targets. Such cell cycle regulated E3 ligases include CRL1^Fbox^ and CRL4^DCAF^, which target many substrates crucial for cell cycle regulation and DNA damage responses (Cardozo and Pagano, 2004; Jackson and Xiong, 2009; Petroski and Deshaies, 2005). These CRLs comprise a scaffolding protein, cullin 1 or cullin 4 (Cul4), an adapter protein (Skp1 and DDB1 respectively), and a RING domain protein that interacts with the E2 (like Rbx1 or Rbx2). Finally, CRL1 and CRL4 ligases contain either an F-box or DCAF substrate recognition factor (SRF, or substrate receptor) respectively, responsible for interacting with the substrate and targeting it for ubiquitination. F-box proteins in CRL1, like Fbw7 or β-TRCP, recognize specific degrons in substrates that often contain phosphorylated residues, while CRL4 include DCAFs like DDB2, which directly recognizes UV-damaged DNA (Scrima et al., 2008).

The CRL4^Cdt2^ ligase uses Cdt2 as the substrate recognition factor, and functions both during S phase and after DNA damage (Abbas and Dutta, 2011; Havens and Walter, 2011; Sakaguchi et al., 2012; Stathopoulou et al., 2012). Cdt2, targets substrates like p21 and Set8, and the DNA replication licensing factor Cdt1 for ubiquitin-mediated proteolysis both in S phase and following DNA damage (Abbas et al., 2008; Centore et al., 2010; Jorgensen et al., 2011; Kim et al., 2008; Nishitani et al., 2008; Oda et al., 2010; Tardat et al., 2010). In addition, an increasing number of Cdt2 target proteins have been identified, including thymine DNA glycosylase (TDG), Cdc6, the DNA polymerase delta subunit p12 (Clijsters and Wolthuis, 2014; Shibata et al., 2014; Slenn et al., 2014; Terai et al., 2013) and Xeroderma pigmentosum group G (XPG), a structure-specific repair endonuclease of the Nucleotide Excision Repair pathway (NER)(Han et al., 2015).

Cdt1 and Cdt2 were originally identified as <u>C</u>dc10 <u>d</u>ependent <u>t</u>ranscript 1 and 2 in fission yeast, but have no sequence similarity (Hofmann and Beach, 1994). Cdt1 has a critical role in establishing the DNA replication licensing complex in G1 phase: it associates with chromatin through the origin recognition complex (ORC) and operates together with Cdc6 to load the MCM2-7 complex onto chromatin, thereby licensing DNA for replication (Bell and Dutta, 2002; Blow and Dutta, 2005; Diffley, 2004; Nishitani and Lygerou, 2004; Symeonidou et al., 2012; Tsakraklides and Bell, 2010).

Preventing re-licensing of replicated regions is essential (Arias and Walter, 2007; Blow and Dutta, 2005). One of the mechanisms to achieve this is by CRL1^Skp2^ and CRL4^Cdt2^ redundantly mediating Cdt1 destruction in higher organisms. CRL1^Skp2^ (also known as SCF^Skp2^) recognizes a phospho-degron motif on Cdt1 that is created at the initiation of S phase by cyclin dependent kinases (CDKs) (Li et al., 2003; Nishitani et al., 2006; Sugimoto et al., 2004). In contrast, CRL4^Cdt2^ recognizes Cdt1 when bound to the proliferating cell nuclear antigen (PCNA) trimer, through a binding motif (PIP box) in its N-terminal end (Arias and Walter, 2006; He et al., 2006; Higa et al., 2006; Jin et al., 2006; Kim and Kipreos, 2007; Nishitani et al., 2006; Ralph et al., 2006; Sansam et al., 2006; Senga et al., 2006). Both initiation of DNA replication and DNA damage trigger PCNA loading onto chromatin and Cdt1 association with PCNA through its PIP box (Arias and Walter, 2006; Havens and Walter, 2009; Raman et al., 2011; Shiomi et al., 2012b). DNA damage-induced degradation of Cdt1 and other substrates appears to facilitate repair (Mansilla et al., 2013; Tanaka et al., 2017; Tsanov et al., 2014)

Cdt2 recruitment onto chromatin is not fully characterized: recruitment through the Cdt1 PIP-box bound to PCNA and a specific basic residue downstream of the Cdt1 PIP box (Havens et al., 2012; Havens and Walter, 2009; Michishita et al., 2011) or independently of Cdt1 (Roukos et al., 2011) has been reported. Following CLR4^Cdt2^ mediated ubiquitination, Cdt1 is degraded, thus blocking further licensing. The N-terminal region of Cdt2 contains a WD40 repeat domain, predicted to form a substrate recognizing propeller structure similar to the one shown for DDB2 (Havens and Walter, 2011; Scrima et al., 2008). In analogy with DDB2 (Fischer et al., 2011), the N-terminal domain of Cdt2 should bind to the DDB1 WD40 repeat domains BPA and BPC on one side. The other side would be expected to recognize the substrate, in analogy to DDB2. Higher eukaryotic Cdt2 proteins have an extended C-terminal region, not present in fission yeast. In *Xenopus* the C-terminal domain of Cdt2 binds to PCNA and is important for the turnover of the Xic1 cyclin kinase inhibitor (Kim et al., 2010). Recently, we reported that Cdt2 mutated at multiple CDK consensus phosphorylation sites co-localized with PCNA throughout S phase even when most of the substrates were degraded, also suggesting that Cdt2 interacts with PCNA independent of its substrates (Nukina et al., 2018).

To understand how Cdt2 recognizes PCNA and localizes CRL4^Cdt2^ activity during S phase and following UV-irradiation, we investigated the role of Cdt2 domains in localization and ubiquitination. Unexpectedly, we found a PIP box in the C-terminal end of Cdt2, and showed that it directly mediates interaction of Cdt2 with PCNA. Both Cdt1 and Cdt2 PIP box peptides bind the PCNA binding pocket in a similar manner, but Cdt2 has significantly higher affinity for PCNA. We suggest that Cdt2 and Cdt1 could simultaneously recognize different subunits of the PCNA trimer, and we put forward a new paradigm for localizing E3 ligase activity onto the PCNA docking platform, allowing simultaneous docking of substrates in the same platform, enabling ubiquitination by proximity.

## RESULTS

### Cdt2 is stably bound to UV-damaged sites independently of Cdt1

CRL4^Cdt2^ recognizes Cdt1 bound on PCNA and recruitment of CRL4^Cdt2^ to damaged chromatin was reported to require Cdt1 in Xenopus extracts(Havens and Walter, 2009). In contrast, experiments in live human cells suggested that Cdt2 was recruited to DNA damage sites independently of Cdt1 (Roukos et al., 2011). To verify whether Cdt2 binding to damaged sites requires Cdt1, we performed a time course analysis of Cdt2 recruitment to sites of localized UV-C irradiation in control and Cdt1 depleted HeLa cells. Cells were exposed to UV-C through micropore filters, and locally induced damage was detected by staining of Cyclopyrimidine dimers (CPDs). In control cells, Cdt1 was rapidly recruited to damaged sites (Fig 1A, 10 minutes) and proteolysed by 30 minutes (fig. 1A and B), consistent with earlier studies (Ishii et al., 2010; Roukos et al., 2011). Cdt2 was recruited at sites of damage by 10 minutes while cells showing Cdt2 recruitment increased at subsequent time points (30 and 45 minutes post-irradiation) even-though Cdt1 was degraded. Depletion of Cdt1 by siRNA had no effect on either the kinetics or the extent of Cdt2 recruitment to sites of UV-C damage (Fig. 1A and 1B). These results show that Cdt2 is recruited to UV-C irradiated sites independently of Cdt1 and remains there long after Cdt1 proteolysis.

**Figure 1.**
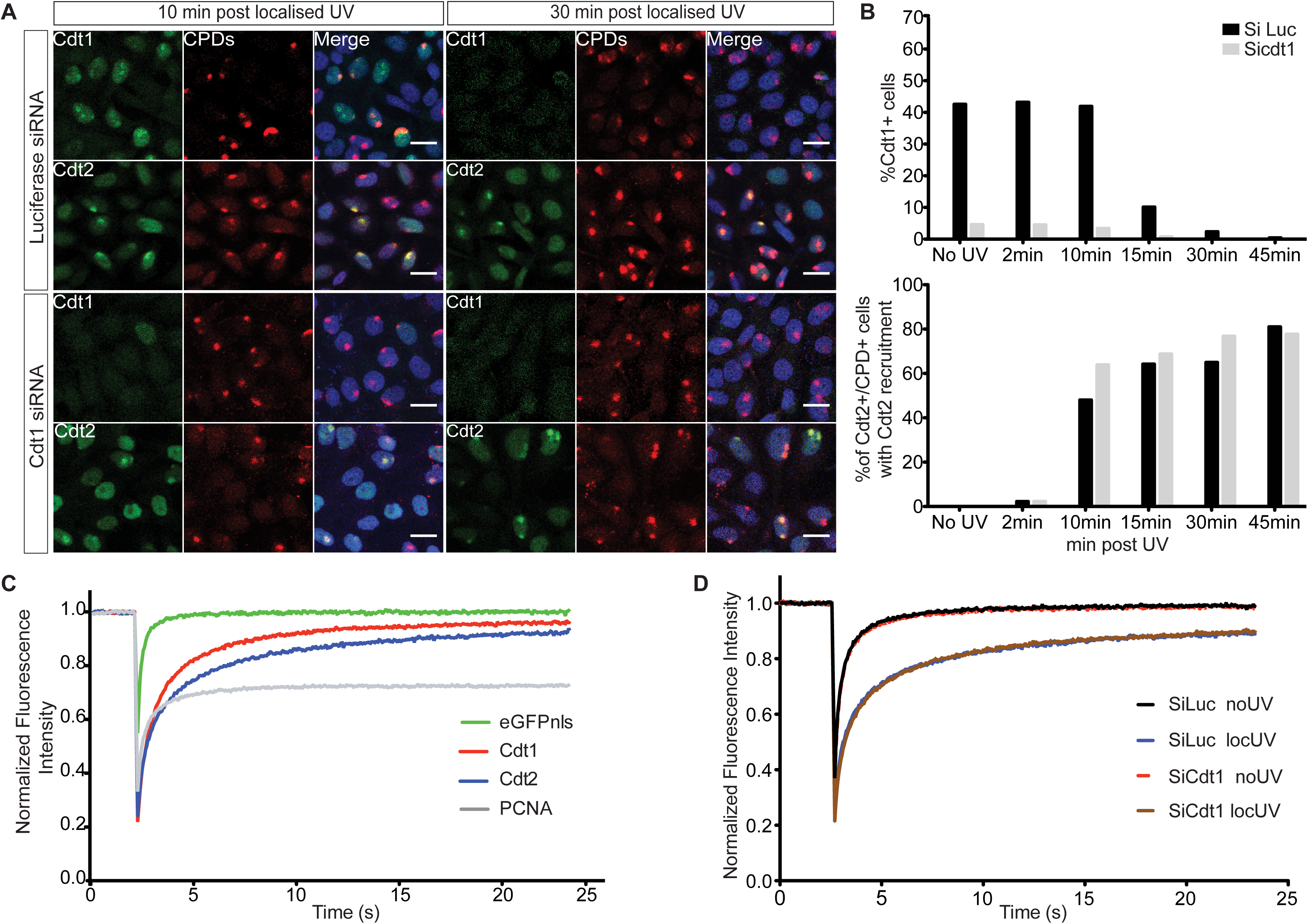
A. HeLa cells were transfected with siRNAs targeted to Cdt1 (siCdt1) or control (siLuc), and locally UV-C irradiated (50 J/m^2^ through a micropore filter with 5μm diameter pores), fixed at the indicated time points and stained with antibodies against CPDs and Cdt1 or Cdt2. Nuclei were stained with Draq5. Scale bars: 20μm. **B**. Percentage of cells positive for Cdt1 (top) and percentage of cells positive for CPDs and Cdt2 which show Cdt2 recruitment to sites of damage (bottom) are plotted. **C.** MCF7 cells were transfected with the indicated GFP-tagged plasmids and 24 hours later they were locally UV-C irradiated (50 J/m^2^). 20 min following irradiation, FRAP experiments were conducted at the site of damage and in untreated cells (Supplementary Fig S1). FRAP data were analysed with easyFRAP. Mean normalised fluorescence intensities at the site of damage as a function of time following photobleaching are shown. **D.** MCF7 cells were synchronised in S-phase by treatment with 2 mM Thymidine for 24 hours. Upon release, cells were transfected with siCdt1 or control siLuc in parallel with a plasmid expressing GFP-tagged Cdt2. Cells were then sunchronised in M-phase by treatment with 50 ng/ml Nocodazole for 12 hours, released into G1 for 5 hours and locally UV-C irradiated (50 J/m^2^). 20 min after irradiation, cell were analysed by FRAP. Mean normalised fluorescence recovery curves at sites of damage are shown.

To assess the binding properties of Cdt2 at sites of UV-C damage in the presence and absence of Cdt1, Fluorescence Recovery after Photobleaching (FRAP) was employed. MCF7 cells transiently expressing Cdt1, Cdt2 or PCNA tagged with GFP were first analysed in the presence and absence of local UV-C irradiation (Fig 1C and Supplementary Fig S1A-C). All factors exhibit fast recovery in the absence of damage (Fig S1A, C), consistent with transient interactions (Xouri et al, 2007b). Following recruitment to sites of damage, PCNA shows a significant immobile fraction, as expected for stable binding, while Cdt1 shows dynamic interactions at UV-C damaged sites (Rapsomaniki et al., 2015; Roukos et al., 2011). Cdt2 has a slower fluorescence recovery rate at the site of damage than Cdt1, with a half-recovery time of 0.83 seconds and an immobile fraction of 16%, underlining long-term association at UV-C damage sites. To assess whether Cdt2 binding to sites of damage is affected by the presence of Cdt1, Cdt2 kinetics were assessed by FRAP in MCF7 cells synchronized in G1, depleted of Cdt1 and locally UV-C irradiated. As shown in Fig 1D and in Supplementary Fig 1D and E, the binding kinetics of Cdt2 at the sites of damage remain the same despite the absence of Cdt1.

We conclude that Cdt2 binds to sites of damage stably independently of Cdt1.

### The C-terminal part of Cdt2 is required for recruitment to DNA damage sites

The above results suggested that Cdt2 contains a domain that mediates its association with UV damaged sites independently of Cdt1. Human Cdt2 is a 730 amino acid polypeptide, with an N-terminal WD40 domain predicted to form a substrate receptor and a long C-terminal domain (Figure 2A). Constructs expressing Cdt2^1-417^, which contains the N-terminal WD40 domain; the C-terminal Cdt2^390-730^; and the full length Cdt2^1-730^ as a control fused with GFP were transiently expressed in MCF7 cells, followed by localised UV-C irradiation (Figure 2). Cdt2^1-730^ was robustly detected at DNA damage sites, as previously reported. The C-terminal Cdt2^390-730^ construct was also efficiently recruited to DNA damage sites. On the contrary, there was no evidence for recruitment of Cdt2^1-417^ to sites of damage. This suggests that the C-terminal region of Cdt2, but not the predicted N-terminal substrate receptor domain, is important for its recruitment to sites of damage.

**Figure 2.**
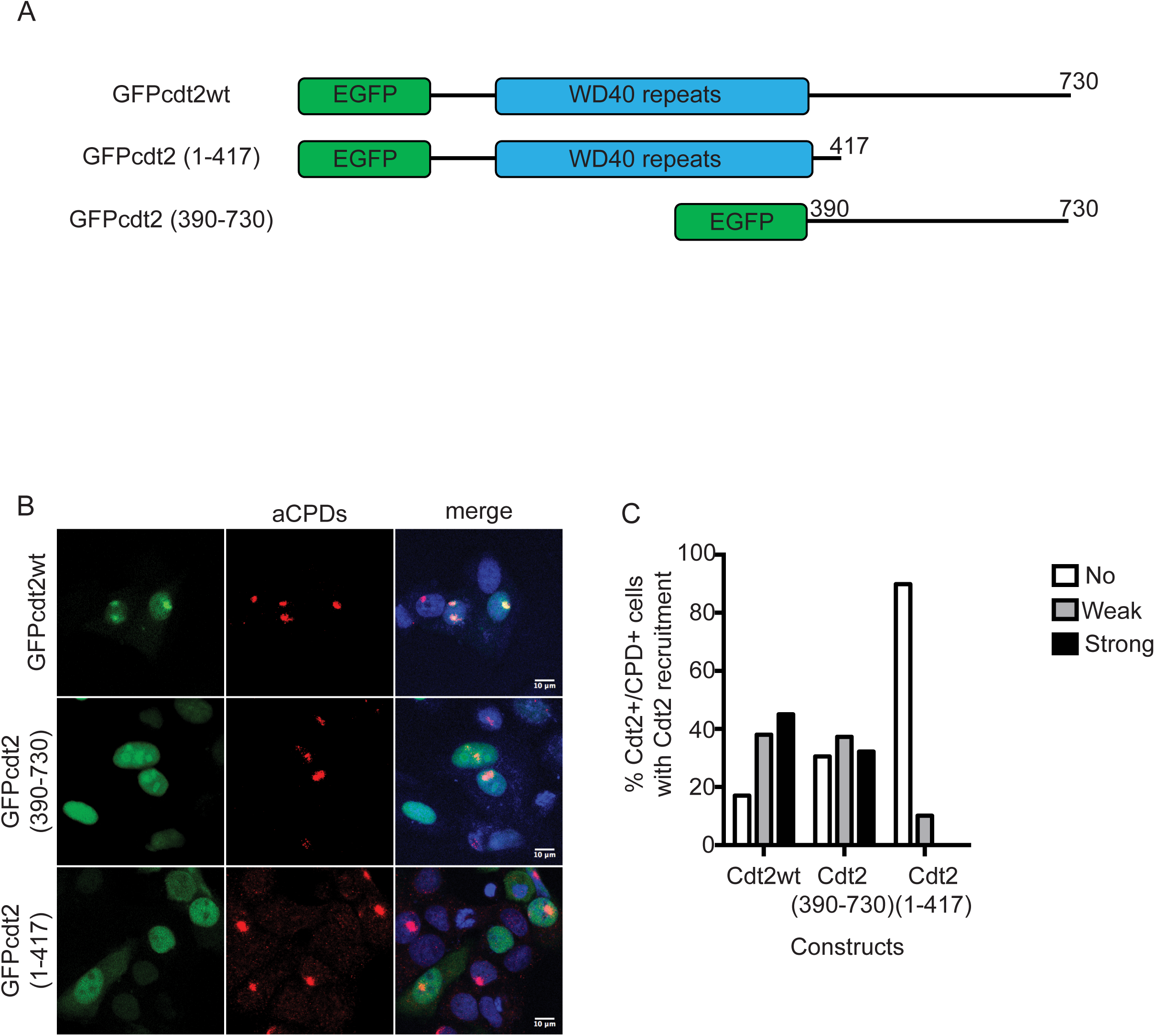
The C-terminus of Cdt2 mediates Cdt2 recruitment to sites of damage. **A.** A schematic drawing of the domain structure of Cdt2 and constructs used. **B** The C-terminal, but not the N-terminal, part of Cdt2 mediates recruitment to DNA damage sites. MCF7 cells transfected with the indicated plasmids were locally UV-C irradiated (50 J/m^2^ through 5μm pore diameter micropore filters), and recruitment of each Cdt2 construct to sites of damage was examined by staining with α-CPD antibody and GFP signal. The percentage of CPD and Cdt2 positive cells which showed strong, weak or no Cdt2 signal at CPD sites is plotted.

### The C-terminal part of Cdt2 is required for interaction with PCNA and has a PIP box in its C-terminal end

Consistent with the observation that the C-terminal region of Cdt2 is important for recruitment to the CPD stained sites where PCNA also localizes, we identified PCNA as a Cdt2 C-terminus interacting protein in a yeast two hybrid screening (Supplementary Fig.S2). To confirm the interaction with PCNA, we transiently transfected 3FLAG-tagged Cdt2^1-730^, Cdt2^1-417^ and Cdt2^390-730^ into HEK293T cells. Before cell lysate preparation, cells were cross-linked. Immuno-precipitation (IP) experiments using α-FLAG antibody showed that the C-terminal half, but not the N-terminal half, interacts with PCNA (Figure 3A). The Cdt2^1-417^ domain, while it lost most of its affinity for PCNA, is still able to bind DDB1 as expected. In contrast, the Cdt2^390-730^ domain, while it still binds to PCNA, has lost its ability to bind DDB1 and therefore to form a functional CRL4 complex.

**Figure 3.**
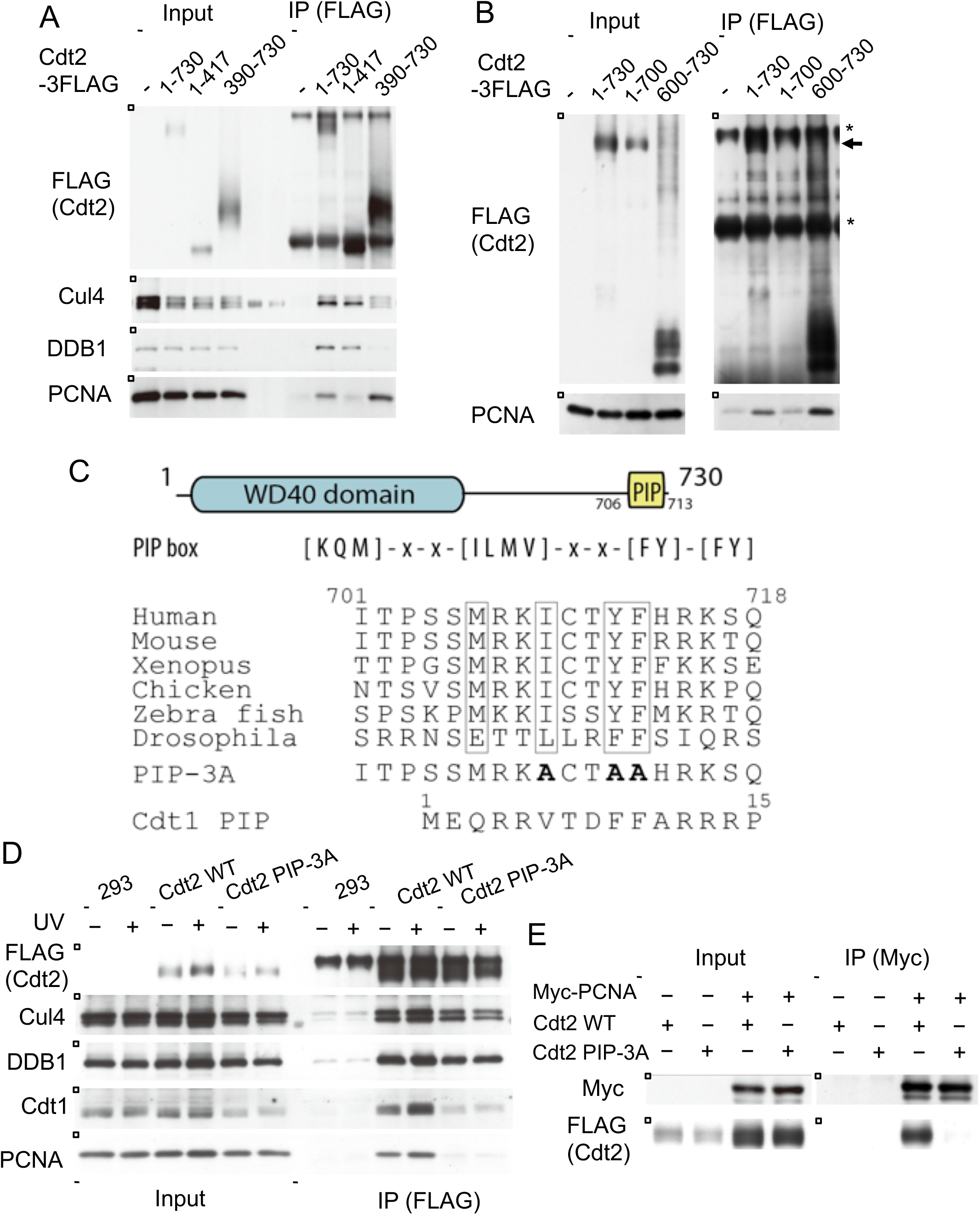
The C-terminal domain of Cdt2 has a conserved PIP-box and interacts with PCNA. **A.** Full length Cdt2 (Cdt2^1-730^) and Cdt2^390-730^, but not Cdt2^1-417^, interact with PCNA. HEK293 cells were transfected with plasmids expressing the indicated Cdt2 fused to a C-terminal FLAG tag, cross-linked and precipitated with anti-FLAG antibodies. Precipitates were examined for PCNA, Cul4A and DDB1. **B**. Cdt2^600-730^, but not Cdt2^2-700^, interact with PCNA. HEK293 cells were transfected with the indicated plasmids and treated as in A. * indicates non-specific bands derived from IgGs. **C**. A schematic drawing of the Cdt2 domain structure highlighting the C-terminal PIP box and a sequence alignment of PIP boxes in various organisms. The alanine changed PIP box mutant of Cdt2 was shown (PIP-3A) together with Cdt1 PIP box. **D**. HEK293 cells were transfected with Cdt2^WT^ or PIP box mutant (Cdt2^PIP-3A^) C-terminally fused to a FLAG-tag, treated as in B and Cdt2 was precipitated with anti-FLAG resin. The interaction with PCNA and DDB1 was examined by immunoblotting. * indicates non-specific bands. **E.** HEK293 cells were transfected with myc-tagged PCNA and Cdt2^WT^-3FLAG or Cdt2^PIP-3A^-3FLAG; PCNA was immunoprecipitated by anti-myc antibodies and the interaction with Cdt2 was examined by immunoblotting.

To narrow the region of Cdt2 required for interaction with PCNA, we made a series of additional constructs fused to a triple FLAG tag. We confirmed that when the C-terminal 30 amino acids were deleted in the Cdt2^1-700^ construct, this was sufficient to lead to loss of interaction with PCNA (Figure 3B). Consistently, the C-terminal 130 amino acids alone, Cdt2^600-730^, were sufficient to mediate the interaction with PCNA.

We postulated that the C-terminal part of Cdt2 directly interacts with PCNA via a PCNA interaction protein (PIP) box; a common motif found in PCNA-dependent DNA replication and repair factors. We used PROSITE to search the UniProtKB protein sequence database for human sequences that contained consensus PIP-boxes ([KMQ]-x-x-[ILMV]-x-x-[FY]-[FY]). Our search revealed a PIP-box sequence from amino acids 706 to 713 in the C-terminus of Cdt2. This motif was well conserved in vertebrates and in *Drosophila* (Figure 3C). We then generated a triple-mutant where the three conserved hydrophobic residues I709, Y712 and F713, also known to interact with PCNA in structures of PIP-box proteins bound to PCNA, were all mutated to alanine, yielding Cdt2^PIP-3A^.

Introducing the Cdt2^PIP-3A^ construct fused to a C-terminal FLAG tag to cells and subsequent IP experiments with anti-FLAG antibody after cross linking, showed that the interaction of the Cdt2^PIP-3A^ construct with PCNA was lost almost entirely (Fig. 3D, UV -), without significantly affecting interaction with Cul4A and DDB1. To confirm the interaction, we performed the reverse IP experiment. Myc-tagged PCNA was co-transfected with Cdt2^WT^ or Cdt2^PIP-3A^ and we performed an IP with anti-myc antibody. Cdt2^WT^ was co-precipitated with myc-PCNA, but Cdt2^PIP-3A^ was not detected in the precipitates (Fig. 3E).

Next, we examined the interaction of CRL4^Cdt2^ ligase with its target protein Cdt1, using HEK293 cells alone and cells transfected with Cdt2^WT^ or Cdt2^PIP-3A^, with or without UV irradiation. Using an anti-FLAG antibody resin to precipitate Cdt2^WT^, both PCNA and Cdt1 were precipitated, and their amounts were notably increased after UV-irradiation (Figure 3D). In contrast, neither Cdt1 nor PCNA co-precipitated well with Cdt2^PIP-3A^, before or after UV irradiation. These results suggested that the PIP-box of Cdt2 was required for forming a stable complex with its substrates on PCNA.

### A C-terminal PIP-box in Cdt2 directly interacts with PCNA with high affinity

To confirm that this PIP box motif in Cdt2 interacts directly with PCNA and to quantify that interaction, we synthesized the peptide corresponding to the human Cdt2 PIP-box (704-717) (SSMRKICTYFHRKS) with a carboxytetramethylrhodamine fluorescent label at the N-terminus (Cdt2^PIP^) and characterized its binding to recombinant PCNA, by fluorescence polarization (FP). Cdt2^PIP^ binds PCNA with high affinity (57±3 nM, Figure 4A). The corresponding Cdt2^PIP-3A^ peptide showed practically no detectable binding to PCNA in the same assay (Figure 4A), suggesting that the Cdt2 PIP-box binds PCNA in a manner very similar to other known PCNA complexes with PIP-box peptides.

**Figure 4.**
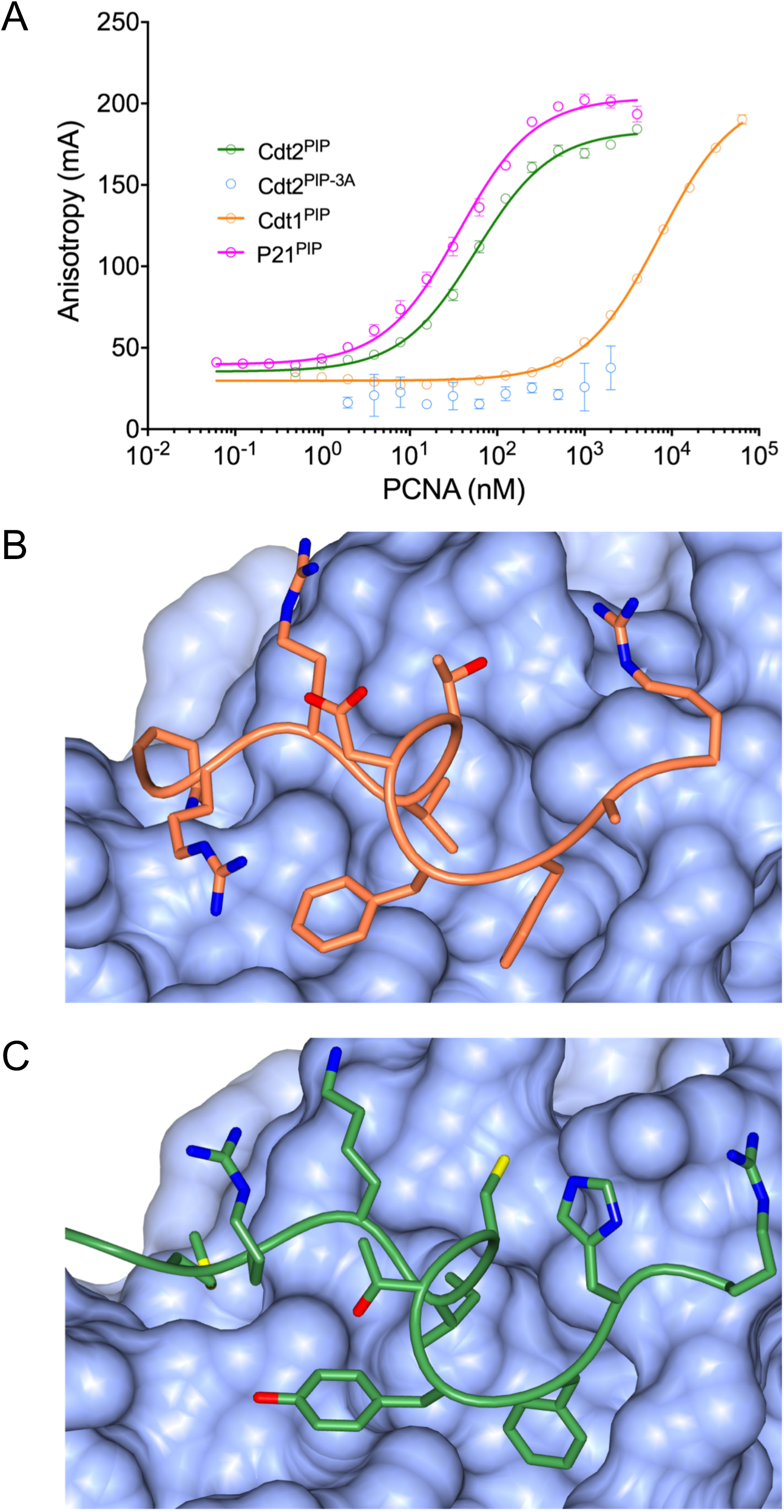
Cdt2 PIP peptide directly interacts with PCNA. **A**. A fluorescence polarization assay showing that the equilibrium binding constants of the Cdt2 PIP-box peptide (Cdt2^PIP^), the triple-mutant of key interacting residues of the classical PIP box (Cdt2^PIP-3A^), as well as the Cdt1 PIP-box peptide (Cdt1^PIP^) and the p21 PIP-box peptide (p21^PIP^). **B.** Crystal structure of the Cdt2^PIP^ peptide bound to PCNA;PCNA is shown as a blue surface; the Cdt2 peptide backbone is shown as an orange “worm” model with side chains as cylinders; carbon atoms are orange; oxygens red and nitrogens blue. **C.** Crystal structure of the Cdt1^PIP^ bound to PCNA; colors as before, but the Cdt1^PIP^ peptide is in green and sulfur yellow.

As Cdt1 is recruited to PCNA on chromatin during S phase and after DNA damage through the Cdt1 N-terminal PIP-box (residues 3-10) (Havens and Walter, 2009; Ishii et al., 2010; Roukos et al., 2011), we then synthesized a Cdt1^PIP^-TAMRA (MEQRRVTDFFARRR) peptide, to compare its affinity to PCNA. Remarkably, the affinity of the Cdt1^PIP^ peptide to PCNA (7200 ± 200 nM, Figure 4A) is two orders of magnitude weaker than that for the Cdt2^PIP^, and similar to what is reported for other peptides derived, for example, from the pol-δ p66 and FEN1 (1-60 µM), (Bruning and Shamoo, 2004). The affinity of the Cdt2^PIP^ peptide for PCNA is similar to that reported for the tightly binding p21 PIP box peptide (50-85 nM) (Bruning and Shamoo, 2004; De Biasio et al., 2012), as it was confirmed in our assays (RRQTSMTDFYHSKR, 36±2 nM, Figure 4A).

To confirm the binding mode of the Cdt2^PIP^ and Cdt1^PIP^ peptides to PCNA, we determined the crystal structure of both complexes by X-ray crystallography, to 3.5 and 3.4 Å resolution respectively. PCNA adopts its well-characterized trimer conformation, with one peptide bound per monomer. The binding mode of both the peptides was very clear (Figure 4B,C, and Suppl. Figure S3A,B, Table 1, and Materials and Methods for details). Both peptides bind similar to other PCNA complex structures. Some notable differences in the binding mode are the conserved Phe713/Phe11 side chain ring that rotates 180 degrees between the two structures and the Tyr712/Phe10 that repositions so as to extend more towards a hydrophilic environment in Cdt2. The Ile709 and Val7 side chains occupy a similar space in the hydrophobic recognition pocket. Albeit the peptide main chain is in a very similar conformation between the structures, some of the other side chains adopt rather different conformations. The average buried area upon the binding of the Cdt2 peptide is 692±6 Å^2^ and average calculated energy of binding is −13±0.2 kcal mol^-1^ while the average buried area upon the binding of the Cdt1 peptide is 650±4 Å^2^ and average calculated energy of binding is −8.3±0.5 kcal mol^-1^; these confirm the tighter binding of the Cdt2 peptide to PCNA.

**Table 1:** data collection and refinement statistics

| PDB identifier | Cdt1 | Cdt2 |
| --- | --- | --- |
| Data Collection |  |  |
| Wavelength (Å) | 1.0000 | 0.8726 |
| Resolution (Å) | 38 – 3.4 (3.63 – 3.40) | 46 – 3.5 (3.83 – 3.50) |
| Space Group | P 2<sub>1</sub> 2<sub>1</sub> 2<sub>1</sub> | P 6<sub>1</sub> |
| Unit Cell a, b, c (Å) | 76.58, 143.73, 173.59 | 151.4, 151.4, 91.49 |
| CC<sub>1/2</sub> | 0.982 (0.647) | 0.976 (0.713) |
| R<sub>merge</sub> | 0.166 (0.631) | 0.289 (0.922) |
| \|I/σ I\| | 7.6 (1.9) | 7.1 (2.1) |
| Completeness (%) | 99.3 (99.2) | 97.1 (88.1) |
| Multiplicity | 3.4 (3.5) | 8.0 (7.1) |
| Unique Reflections | 26,761 | 14, 830 |
| Refinement |  |  |
| PCNA/Cdt2 peptides in A.U. | 6 | 3 |
| Atoms protein | 24,564 | 12,324 |
| B-factors protein (Å<sup>2</sup>) | 48.0 | 45.8 |
| R<sub>work</sub>/R<sub>free</sub> (%) | 20.5/25.1 | 20.0/25.6 |
| Bond lengths rmsZ/rmsd (Å) | 0.0087/0.612 | 0.083/0.600 |
| Bond angles rmsZ/rmsd (°) | 1.682/0.981 | 1.708/0.999 |
| Ramachandran preferred/outliers (%) | 89.1/1.9 | 87.8/1.7 |
| Clash score (%-ile) | 97 | 97 |
| MolProbity score (%-ile) | 92 | 90 |

High Resolution shell in parentheses

### Cdt2 directly binds to PCNA on DNA through its PIP box independently of Cdt1 *in vitro*

The above results suggested that Cdt2 was recruited to PCNA sites primarily through its PIP-box rather than the N-terminal substrate receptor domain, which contrasts to the model that Cdt1 and CRL4^Cdt2^ are sequentially recruited to PCNA^on Chromatin^. To investigate the mechanism of Cdt1 and CRL4^Cdt2^ recruitment to PCNA, we set up an *in vitro* analysis with purified and defined human proteins. FLAG tagged Cdt1(Cdt1-3FLAG) and CRL4^Cdt2^ (Cdt2-3FLAG) were expressed in insect cells and purified on anti-FLAG resin (Figure 5A). PCNA and its loader RFC were purified as described in the Experimental procedures (Supplementary Figure S4). First, we examined the interaction of Cdt1 or CRL4^Cdt2^ with PCNA in the absence of DNA (free PCNA). The purified Cdt1 or CRL4^Cdt2^ were fixed on anti-FLAG beads, incubated with free PCNA, and the amount of bound PCNA was analyzed. In contrast to the above mentioned results that both Cdt1 and Cdt2 PIP-box peptides bound to PCNA (Figure 4), PCNA was detected on Cdt1-beads, but much less on CRL4^Cdt2^-beads (Figure 5B).

**Figure 5.**
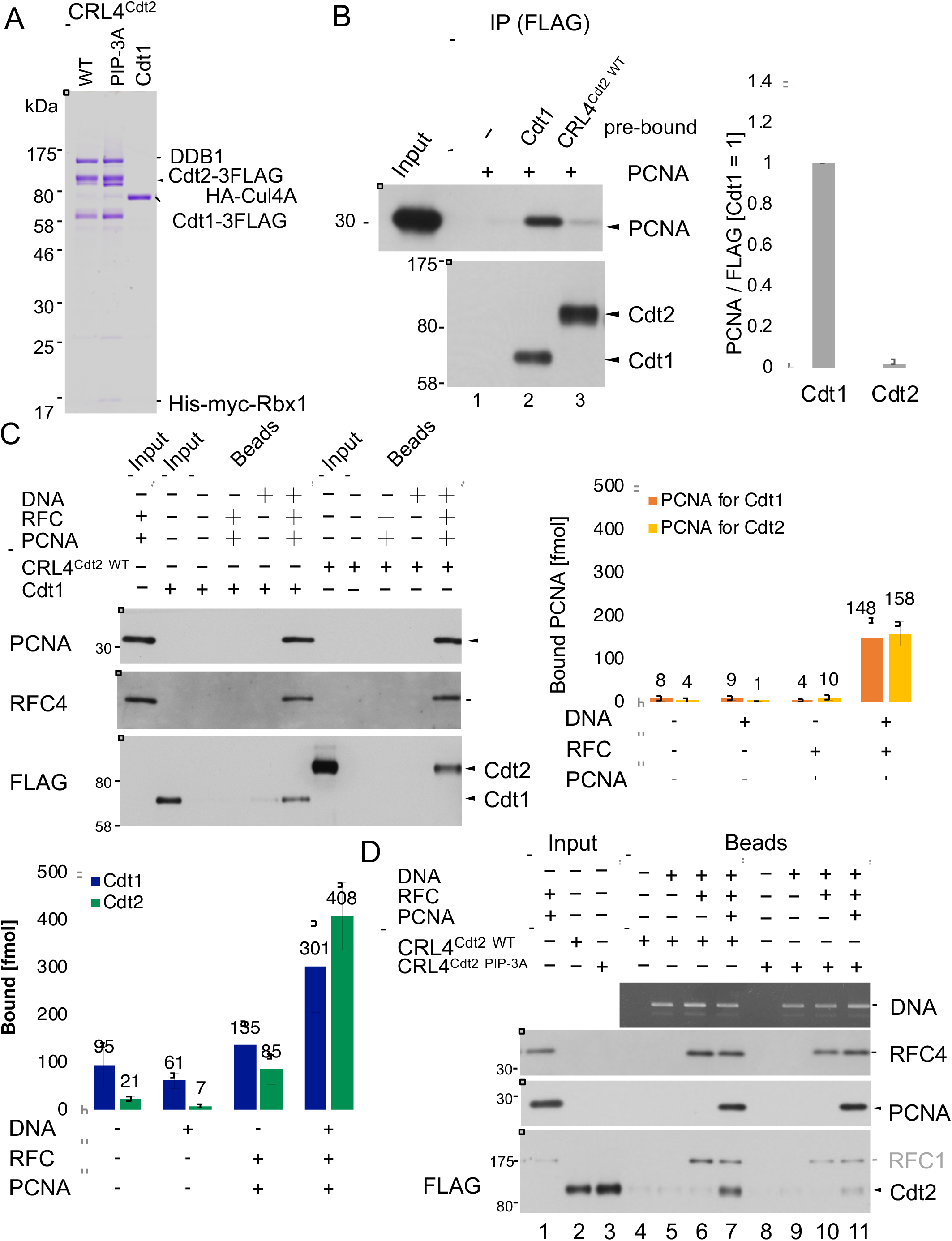
Cdt1 and CRL4^Cdt2^ independently associate with PCNA loaded on DNA through PIP-box *in vitro*. **A**. Purified CRL4^Cdt2(WT)^, CRL4^Cdt2(PIP-3A)^ and Cdt1(WT) proteins. **B**. Cdt1, but not CRL4^Cdt2^, associates with free PCNA. The purified Cdt1 and CRL4^Cdt2^ proteins were fixed on anti-FLAG-resins and incubated with PCNA for 2hrs at 4 °C and the bound PCNA was analyzed. The relative amounts of bound PCNA were shown, normalized with FLAG signal and set the level on Cdt1-3FLAG beads to 1.0. **C**. The ncDNA-beads (DNA+) or control beads (DNA-) were loaded with PCNA by RFC or not. Then, beads were incubated with the purified Cdt1 or CRL4^Cdt2^, and the bound proteins were recovered for Western blotting. The amounts of PCNA loaded on plasmid DNA and the amounts of bound Cdt1-3FLAG and Cdt2-3FLAG were measured and shown (f mol). **D**. CRL4^Cdt2(WT)^ and CRL4^Cdt2(PIP-3A)^ were incubated with control beads, ncDNA beads or ncDNA beads which had been pre-incubated with RFC alone or together with PCNA for 1 hr at 4 °C. Beads were recovered and bound proteins were analysed.

Because Cdt1 and Cdt2 bind to PCNA only when PCNA is loaded on DNA in *Xenopus* egg extracts (Havens and Walter, 2009), we prepared PCNA loaded on nicked circular DNA (ncDNA) with RFC (PCNA^on DNA^) (Supplementary Figure S5). In our *in vitro* conditions, three molecules of PCNA trimer were loaded on one plasmid DNA. Then, we analyzed the binding of Cdt1 and CRL4^Cdt2^ to PCNA^on DNA^ as described in Supplementary Figure S5A. After incubation, Cdt1 was efficiently recovered on the PCNA^on DNA^ -beads, but not on the control beads (Figure 5C). Although we could not show CRL4^Cdt2^ binding to free PCNA, we show that CRL4^Cdt2^ binds to PCNA^on DNA^ in the absence of Cdt1. The interaction was not mediated by RFC proteins bound on ncDNA (Figure 5D lane 6). The molar ratio of Cdt2 bound to trimeric PCNA^on DNA^ was the same or somewhat higher than that of Cdt1 (406 fmol of Cdt2 bound to 158 fmol trimeric PCNA^on DNA^, while 301 fmol of Cdt1 to 148 fmol of trimeric PCNA^on DNA^). These results indicate that CRL4^Cdt2^ directly and efficiently binds to DNA-loaded PCNA without substrate, consistent with data in cells showing that Cdt2 was recruited to sites of damage in the absence of Cdt1 (Figure 1).

To confirm that the interaction of CRL4^Cdt2^ with PCNA^on DNA^ was mediated by the PIP-box of Cdt2, the CRL4 complex having Cdt2^PIP-3A^ (CRL4^Cdt2(PIP-3A)^) was purified like CRL4^Cdt2^ (Figure 5A) and used in a binding assay. As above, CRL4^Cdt2^ interacted with PCNA^on DNA^. In contrast, the binding activity of CRL4^Cdt2(PIP-3A)^ to PCNA^on DNA^ was severely reduced as compared with CRL4^Cdt2^ (Figure 5D). These results demonstrate that the PIP-box of Cdt2 was directly responsible for the interaction of CRL4^Cdt2^ with PCNA^on DNA^.

### The Cdt2 PIP-box promotes Cdt2 recruitment to UV irradiated sites and PCNA foci

To investigate the role of the Cdt2 PIP-box in cells we isolated HEK293 cells stably expressing FLAG tagged Cdt2^PIP-3A^ and showed that Cdt2^PIP-3A^ was not recruited to UV-irradiated sites (Figure 6A). XPA, a component of nucleotide excision repair, staining was used to confirm that similar extents of DNA damage were induced both in Cdt2^WT^ and Cdt2^PIP-3A^ expressing cells. Next, we verified that the recruitment of Cdt2 onto PCNA during S phase is also dependent on its PIP-box. In order to detect the chromatin-associated fraction of PCNA and Cdt2, asynchronous cells were pre-extracted prior fixation and co-stained with PCNA and FLAG antibodies. A similar percentage of PCNA-positive S phase cells was detected both in Cdt2^1-730/WT^ expressing cells and Cdt2^PIP-3A^ expressing cells (Supplementary Figure S6). Cdt2^WT^ was co-localized with PCNA foci, corresponding to sites of DNA replication. More than 80% of PCNA-positive cells were co-stained with Cdt2^WT^, with the early S phase cells displaying stronger Cdt2^WT^ staining than the late S phase cells as reported (Nukina et al., 2018) (Figure 6B and 6C and Supplementary Figure S6A). In contrast, almost no signal was detected for Cdt2^PIP-3A^ irrespective of the presence of PCNA. Consistent with the immunofluorescence analysis, following cell fractionation, Cdt2^WT^ was recovered in a chromatin-containing fraction, while Cdt2^PIP-3A^ was undetectable (Figure 6D).

**Figure 6.**
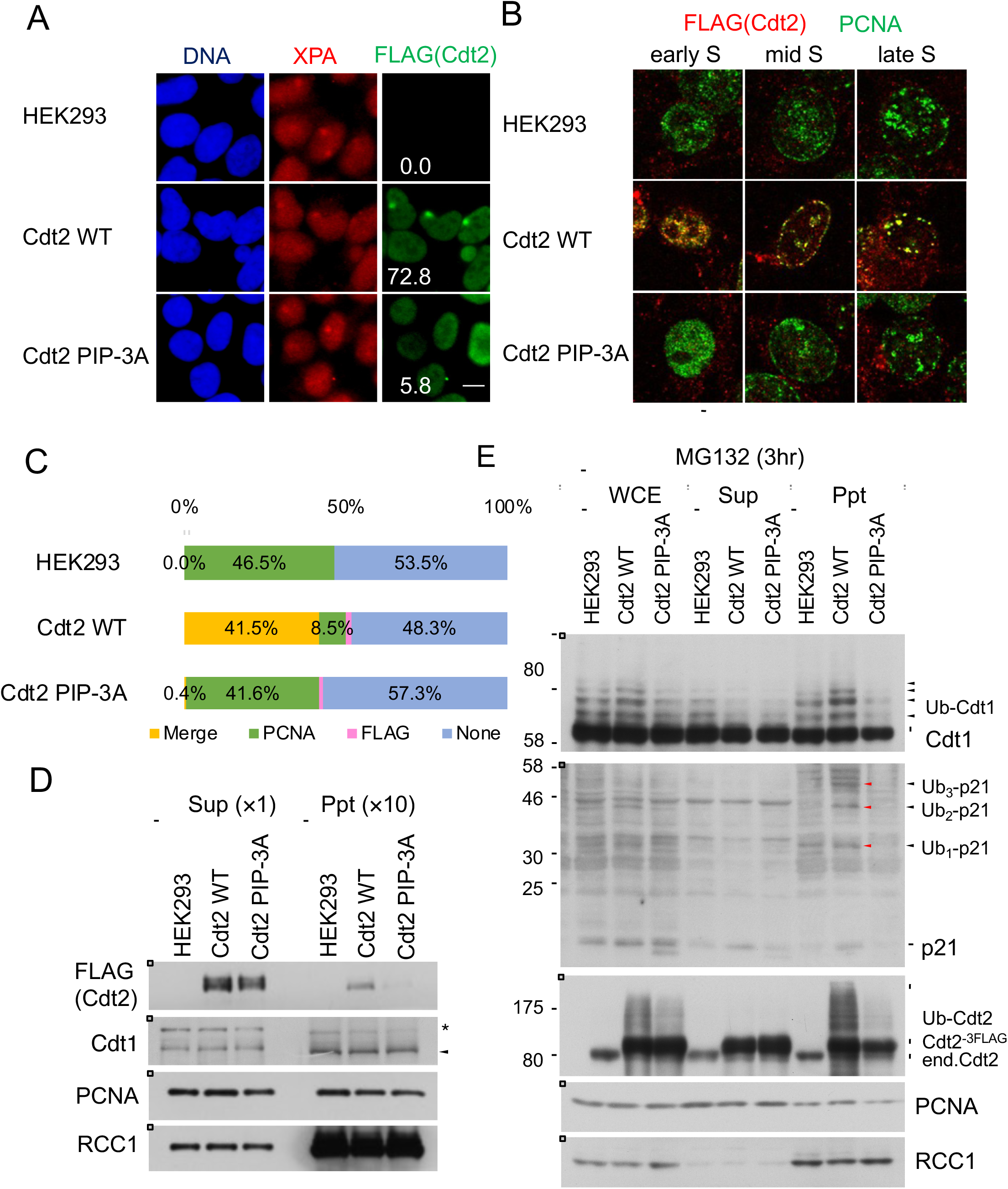
The Cdt2 PIP-box is important for its recruitment to DNA damage sites and replication foci and for poly-ubiquitination activity. **A**. HEK293 and stable Cdt2WT or Cdt2PIP-3A cells were locally UV-irradiated, and stained for XPA and FLAG (Cdt2). XPA-positive spots were examined for FLAG signal, and its frequency was counted (%). More than 50 spots were examined. bar:10 μm. **B**. HEK293 cells and stable cells were pre-extracted, fixed and stained with PCNA and DYKDDDDK (FLAG) antibodies. bar:10 μm. **C**. Cells stained as (B) were examined for staining with PCNA and DYKDDDDK (FLAG) antibodies and the frequency of cells with colocalized PCNA and FLAG (Cdt2) staining (Merge), PCNA staining alone (PCNA), FLAG staining alone (FLAG) or no staining (None) were plotted. **D**. Whole cell extracts were prepared from HEK293 and stably transfected cells of Cdt2^WT^ or Cdt2^PIP-3A^ cells, separated into soluble (Sup) and chromatin-containing insoluble (Ppt) fractions and were examined for indicated proteins. **E**. HEK293 and stably transfected cells of Cdt2^WT^ or Cdt2^PIP-3A^ were treated with MG132 for 3 hr. Whole cell extracts (WCE) were prepared, separated into soluble (Sup) and insoluble (Ppt) fractions and blotted with the indicated antibodies.

These results collectively suggest that while Cdt2^PIP-3A^ is capable of forming a complex with the rest of the components of the CRL4 ligase, the Cdt2 PIP-box is required for efficient recruitment of CRL4^Cdt2^ to the PCNA-loaded sites.

### Cdt1 ubiquitination is abortive in Cdt2^PIP-3A^ expressing cells

Since the recruitment of Cdt2^PIP-3A^ to PCNA sites was compromised, it would imply that ubiquitination of Cdt1 was also abrogated. To consider this possibility, asynchronously growing control HEK293 cells and cells expressing Cdt2^WT^ or Cdt2^PIP-3A^ were treated with proteasome inhibitor MG132 and the levels of poly-ubiquitinated Cdt1 were examined. While poly-ubiquitinated Cdt1 was clearly detected in Cdt2^WT^ expressing cells, its levels were reduced in Cdt2^PIP-3A^ expressing cells (Figure 6E), indicating that the ability of Cdt2^PIP-3A^ to ubiquitinate Cdt1 was compromised. Fractionation of cell extracts revealed that most of the poly-ubiquitinated Cdt1 and p21 were recovered in the chromatin-containing precipitate fraction prepared from Cdt2^WT^ expressing cells, in agreement with CRL4^Cdt2^ operating on chromatin (Figure 6E). Cdt2^PIP-3A^ expressing cells however had reduced level of both poly-ubiquitinated Cdt1 and p21 on chromatin. Interestingly, Cdt2 itself is poly-ubiquitinated on chromatin, and this is severely reduced in Cdt2^PIP-3A^ expressing cells (Figure 6E). Increased levels of poly-ubiquitination of Cdt1 and p21 were also observed in another cell line, U2OS cells expressing Cdt2^WT^, but not in cells expressing Cdt2^PIP-3A^ (Supplementary Figure S7B).

Next, we examined the accumulation of poly-ubiquitinated Cdt1 after UV-irradiation. For this experiment, the endogenous Cdt2 was depleted with an siRNA targeted to the 3’ UTR to assay for the activity of introduced Cdt2. Cells were treated with MG132, irradiated with UV and the levels of poly-ubiquitinated Cdt1 were monitored by Western blotting. In HEK293 cells treated with control siRNA, poly-ubiquitinated Cdt1 was detected 15 min after irradiation (Supplementary Figure S6B). Depletion of Cdt2 blocked the poly-ubiquitination of Cdt1. The defect was rescued by Cdt2^WT^ expression. In contrast, Cdt2^PIP-3A^ expressing cells were defective in poly-ubiquitination of Cdt1 (Supplementary Figure S6B), suggesting that Cdt2 PIP-box is required for efficient poly-ubiquitination of substrates.

### The Cdt2 PIP-box is required for efficient Cdt1 degradation

Since the poly-ubiquitination activity of Cdt2^PIP-3A^ was reduced, we monitored the degradation of Cdt1 after UV irradiation. Asynchronously growing control U2OS cells and cells expressing Cdt2^WT^ or Cdt2^PIP-3A^ (Supplementary Figure S7) were depleted of endogenous Cdt2 using an siRNA targeted to the 3’ UTR, and irradiated with UV. When cells were exposed to UV (20 J/m^2^), degradation of Cdt1 was almost prevented in U2OS cell depleted of endogenous Cdt2 (Figure 7A and 7B). Ectopic expression of Cdt2^WT^ restored Cdt1 degradation to similar kinetics to that of control siRNA transfected U2OS cells. In contrast, Cdt2^PIP-3A^ was much less effective in rescuing Cdt2 depletion. Similarly, at a lower dose of UV-irradiation (5 J/m^2^), Cdt1 remained stable 1 hr post UV-irradiation in Cdt2^PIP-3A^ cells, when endogenous Cdt2 was depleted, both on immunofluorescent analyses and Western blotting (Figure 7C and Supplementary Figure S7C).

**Figure 7.**
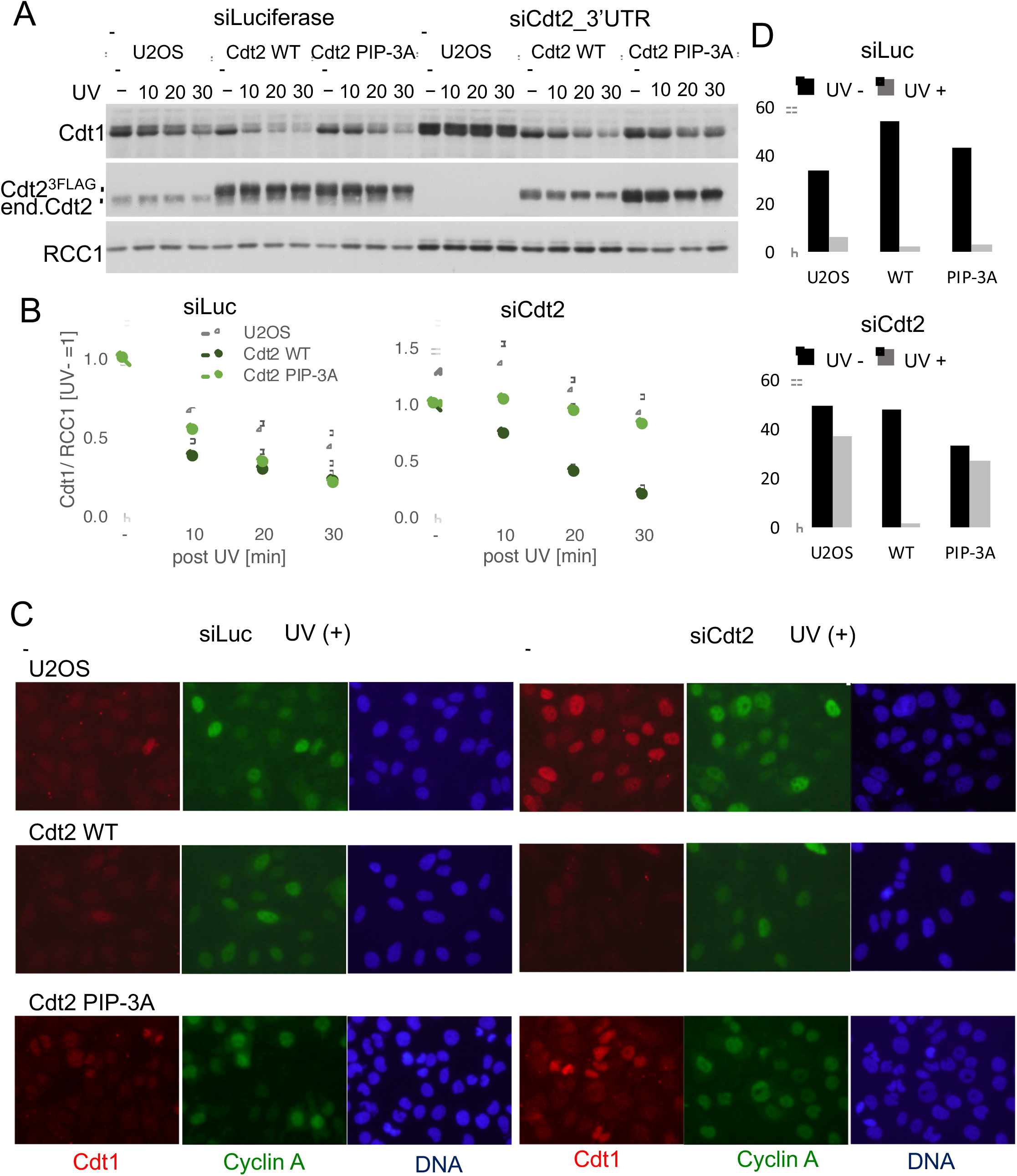
The Cdt2 PIP-box is important role for rapid Cdt1 degradation after UV-irradiation. **A**. U2OS cells and U2OS cells stably expressing Cdt2^WT^-3FLAG or Cdt2^PIP-3A^-3FLAG were transfected with siCdt2 targeted to the 3’ UTR or control siLuc for 48 hr, irradiated with UV (+, 20 J/m^2^) or not (-) and collected at the indicated time points for Western blotting. **B**. The protein levels were measured and the relative amounts of Cdt1, normalized to RCC1, after UV irradaition were shown, set the UV (-) level as 1.0. **C**. Cells were transfected with siCdt2 or control siLuc for 72 hr, irradiated with UV (+, 5 J/m^2^) or not (-) and 1hr later fixed for staining with Cdt1 and CyclinA antibodies. **D**. The relative frequency of Cdt1-positive cells in (C) was shown (%) (average of two independent experiments).

**Figure 8.**
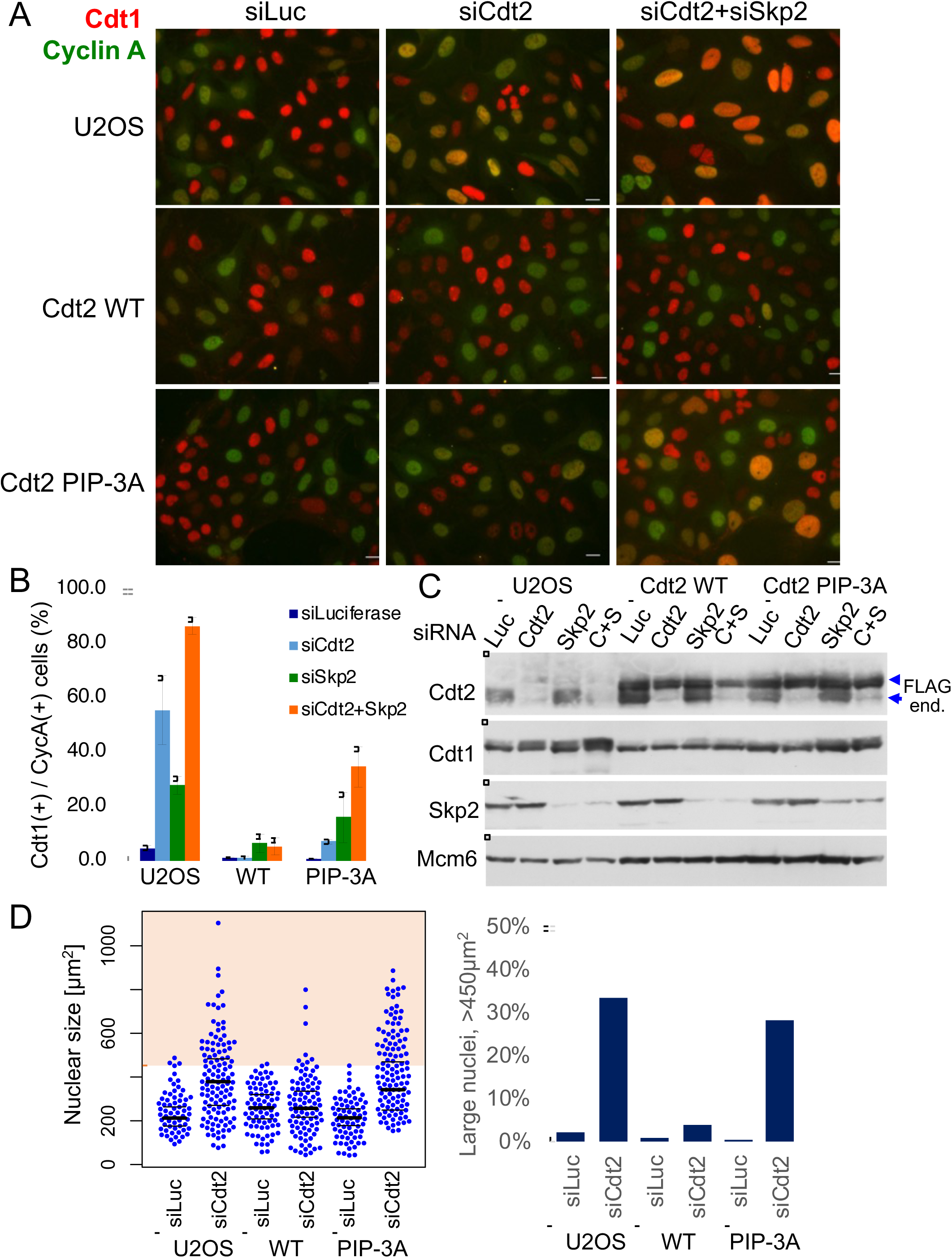
The Cdt2 PIP-box plays an important role for Cdt1 degradation during cell cycle. **A** and **B**. U2OS cells and U2OS cells stably expressing Cdt2^WT^-3FLAG or Cdt2^PIP-3A^-3FLAG were transfected with siCdt2 and/or siSkp2 or control siLuc for 72 hr. Cells were fixed and stained with Cdt1 and CyclinA antibodies. The frequency of Cdt1 positive (+) cells among Cyclin A positive (+) cells was calculated (n=3). **C**. Cells transfected with siRNAs as above (C+S; siRNAs for Cdt2 and Skp2) were examined for indicated protein levels. **D**. Cells treated with siCdt2 or control siLuc were measured for their nuclear size (n=200 for each), and were shown in scatter plot. Frequency of cells with nuclear size larger than 450 μm^2^ is shown (%).

Next, we examined Cdt1 degradation after the onset of S phase in the Cdt2^PIP-3A^ expressing cells. Cyclin A is present in S and G2 phase, when Cdt1 should be degraded. Thus, Cdt1 should be present in cells that are negative for Cyclin A, and absent in cells that are positive for Cyclin A (Nishitani et al., 2006; Xouri et al., 2007a; Xouri et al., 2007b). Cdt1 degradation after the onset of S phase is carried out by two ubiquitin ligases, CRL1^Skp2^ and CRL4^Cdt2^. We depleted the substrate recognition subunits, Cdt2 and/or Skp2, by siRNA, and the percentage of Cdt1 positive cells among Cyclin A-positive cells, was assessed by immunofluorescence. In control siRNA transfected U2OS cells, ~2% of Cyclin A positive cells stained positive for Cdt1. In contrast, in Cdt2 depleted, Skp2 depleted and Cdt2/Skp2 co-depleted cells, 50%, 30% and more than 80% of Cyclin A positive cells stained positive for Cdt1 respectively (Figure 8A-8C), consistent with earlier work (Nishitani et al., 2006). Then, we depleted Cdt2 and Skp2 and examined Cdt1 degradation in Cdt2^WT^ or Cdt2^PIP-3A^ expressing cells. Ectopic expression of Cdt2^WT^ fully complemented the depletion of Cdt2, and also the co-depletion of Skp2. This was probably because the levels of ectopically expressed Cdt2^WT^ were higher than the endogenous Cdt2 levels. On the other hand, Cdt1 degradation was not fully rescued by the expression of Cdt2^PIP-3A^ protein both in Cdt2 depleted and Cdt2 and Skp2 co-depleted cells (Figure 8A and 8B). Upon depletion of endogenous Cdt2, many cells exhibited significantly enlarged nuclei, consistent with rereplication. This phenotype was rescued by ectopic expression of Cdt2^WT^ but not of Cdt2^PIP-3A^ (Fig.8D and Supplementary Fig.S8), suggesting that the PIP-box of Cdt2 is essential to prevent rereplication.

Taken together, our *in vitro* and *in vivo* analyses show that the Cdt2 PIP-box brings CRL4 ubiquitin ligase to PCNA sites and is required for efficient substrate degradation both during S phase and following UV irradiation, to maintain genome stability.

## DISCUSSION

Cdt2 substrates are degraded rapidly as cells enter S phase or when cells are exposed to DNA damaging agents. Cdt2 of higher eukaryotic cells has an extended C-terminal region. We demonstrate that Cdt2 has a PIP box in its C-terminal end that recognises chromatin bound PCNA with high affinity. This interaction is important for CRL4^Cdt2^ function in recognising its targets prior to their ubiquitin-dependent degradation: this previously un-noticed Cdt2 PIP-box is required for recruitment of CRL4^Cdt2^ to chromatin bound PCNA, and indispensable for efficient Cdt1 ubiquitination, thus for prompt degradation of Cdt1 both in S phase and after UV irradiation. The Cdt2 PIP-box peptide directly interacts with PCNA with high affinity (about 50 nM) *in vitro,* similar to the unusually high-affinity p21 PIP box peptide, and significantly tighter than the Cdt1 PIP-box peptide. Molecular modelling experiments suggest that both the Cdt1 and Cdt2 PIP-boxes bind PCNA in the classical way and their binding to PCNA occurs independently of each other (Figure 4). Furthermore, the purified CRL4^Cdt2^ ligase interacted directly to the PCNA loaded on DNA via Cdt2 PIP-box (Figure 5). Consistently, the C-terminal domain of Cdt2 has been shown to bind to PCNA in *Xenopus* egg extracts (Kim et al., 2010). Our analysis now shows that the PIP-box is conserved in *Xenopus,* and thus likely to be responsible for this interaction.

PCNA is a trimer, hence it offers three PIP-box binding sites that are proposed to be asymmetric when PCNA is loaded onto DNA (Ivanov et al., 2006). Consequently, both Cdt1 and Cdt2 can be simultaneously recruited onto PCNA, effectively utilizing PCNA as a common docking platform for bringing the CRL4^Cdt2^ ligase and the Cdt1 substrate in proximity, allowing more efficient transfer of a ubiquitin moiety from the E2 to Cdt1 (Figure 9). This Cdt2 PIP-box assisted ubiquitination is likely to be used by other Cdt2 substrates, as p21 poly-ubiquitination was also defective in Cdt2^PIP-3A^ expressing cells (Figure 6E and Supplementary Figure S7B). This mechanism, where co-localization onto the same protein platform brings the E3 ubiquitin ligase and its substrate in proximity, is a new paradigm in ubiquitin conjugation. This model fits recent results which show that Cdt2 itself can be auto-ubiquitinated by CRL4^Cdt2^ (Abbas et al., 2013): CRL4^Cdt2^ and Cdt2 (or a second CRL4^Cdt2^) may both bind PCNA through the Cdt2 PIP-box and catalyse this auto-ubiquitination event. Consistently, the poly-ubiquitination of Cdt2 which was observed on chromatin in Cdt2^WT^ expressing HEK293 cells was severely reduced in Cdt2^PIP-3A^ expressing cells (Figure 6E).

**Figure 9.**
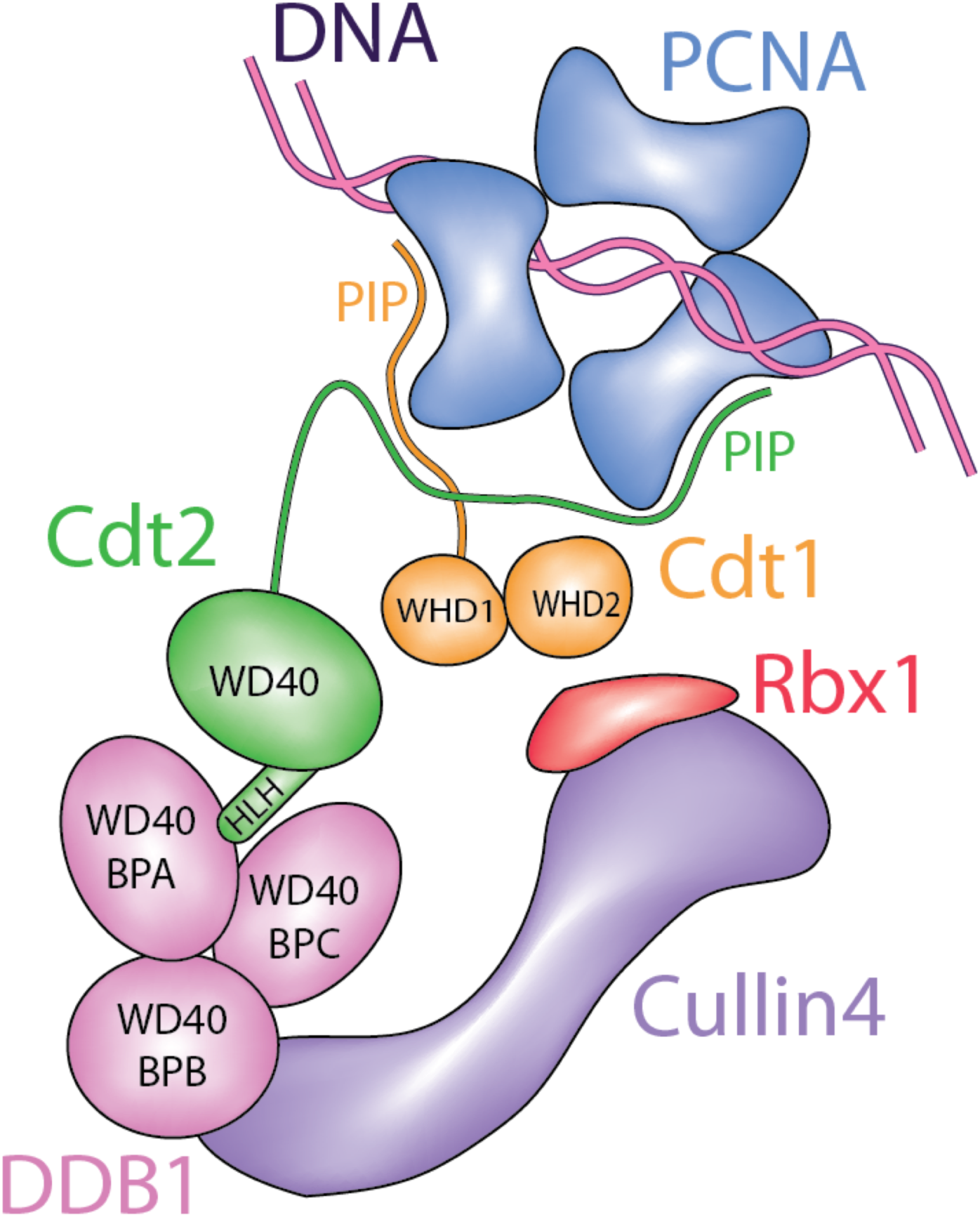
Model. A model showing how both CRL4^Cdt2^ and Cdt1 are likely recruited to the chromatin-bound PCNA through their PIP boxes, for efficient ubiquitination.

The mono-ubiquitination (mUb) of PCNA is carried out by Rad6-Rad18, but also CRL4^Cdt2^ contributes to it. In non-damaged cells, mUb of PCNA is present at a basal level and its levels increase in response to DNA damage (Terai, et al). The Cdt2 PIP-box may also be important for mUb of PCNA, as mUb-PCNA levels appeared decreased in Cdt2^PIP-3A^ in comparison to Cdt2^wt^ expressing cells (Supplementary Figures S7B). Therefore, Cdt2 PIP-box may be required for various events driven by CRL4^Cdt2^ in the cell; degradation of PIP-degron proteins, auto-ubiquitination of Cdt2 and PCNA mono-ubiquitination.

We show that the Cdt2 PIP-box is crucial for recruitment of the CRL4^Cdt2^ ligase onto PCNA in human cells. However, further interactions of Cdt2 domains, mediated by its N-terminal WD40 repeats, with the PCNA-bound Cdt1 PIP-box (the PIP degron) also enhance the affinity of the CRL4^Cdt2^ (Havens and Walter, 2009). Here, we propose a synergistic mechanism where the Cdt2 PIP-box and the Cdt1 PIP-degron operate in concert to promote CRL4^Cdt2^-dependent target ubiquitination (Figure 9). It is also noteworthy that the binding of Cdt1 onto PCNA is transient in cells, while Cdt2 interactions are more stable (Roukos et al., 2011 and Figure 1B): a synergistic mechanism would imply that interactions of pre-bound Cdt2 with transiently formed Cdt1 PIP-degron, are important for localising CRL4^Cdt2^ and Cdt1 onto PCNA.

Although the Cdt2 PIP-box mediates direct interaction with PCNA^on DNA^, the affinity of Cdt2 to PCNA decreases as cells progress into and complete S phase. The chromatin levels of Cdt2 are reduced during late S phase (Rizzardi et al., 2015), and the co-localization of Cdt2^WT^ with PCNA was mostly observed in early S phase cell nuclei (Figure 6B). It was suggested that the CDK-dependent phosphorylation of Cdt2 down-regulates Cdt2 interaction with PCNA, as Cdk1 inhibition recovered the chromatin-interaction of Cdt2 at late S phase, and Cdt2 mutated at CDK phosphorylation sites displayed strong interaction with PCNA throughout S phase (Nukina et al., 2018). We propose that the C-terminal region of Cdt2 contributes to regulating the activity of CRL4^Cdt2^ in the cell cycle by controlling its affinity to PCNA; Cdt2 PIP-box mediated binding to PCNA is important for rapid substrate degradation at the onset of S phase, while the phosphorylation of the C-terminal region of Cdt2 inhibits its PCNA interaction and thus reduces ubiquitination activity during late S phase, leading to substrate re-accumulation. It remains, however, to be elucidated how phosphorylation of Cdt2 inhibits its interaction to PCNA.

CRL-type ligases are thought to use the ‘substrate recognition factor’ or ‘substrate receptor’ subunit to recognize a specific degron in the substrate itself. However, CRL4-type ligases can deviate from that mechanism. For example, CRL4^DDB2^ recognizes specific UV-damaged bases in DNA, creating a ‘ubiquitination zone’ around the repair site, to ubiquitinate specific substrates (Fischer et al., 2011). Here we show a novel specific mechanism, where ligase and substrate recognize the same molecule of a cellular platform, to bring enzyme and substrate into close proximity and thus to facilitate substrate trapping: our data constitute a new paradigm for the use of a docking platform in ubiquitin conjugation. Creating such a three-component system offers many possibilities for regulation and robustness of the ubiquitination response: the recognition of PCNA by both substrate and Cdt2 through specialized PIP boxes likely enables further interactions between substrate and Cdt2 (as for example the recognition of the PIP degron by its N-terminal WD40 repeat domain), creating a set of redundant molecular recognition events that regulate this system in the cellular context.

## EXPERIMENTAL PROCEDURES

### Cell Culture

HeLa cells, HEK293 cells, 293T cells and U2OS cells were cultured in Dulbecco’s modified Eagle’s medium with 10% fetal bovine serum and 5% CO_2_. MCF7 cells were cultured in Dulbecco’s modified Eagle’s medium with 20% fetal bovine serum and 5% CO_2_ at 37°C. HEK293 cells and U2OS cells stably expressing Cdt2^WT(1-730)^-3FLAG, Cdt2^1-417^-3FLAG, FLAG-Cdt2^300-730^ or Cdt2^PIP-3A^-FLAG were isolated using plasmids pCMV-HA-Cdt2^WT(1-730)^-3FLAG, pCMV-HA-Cdt2^1-417^-3FLAG, p3FLAG-Cdt2^300-730^-3NLS-myc or pCMV-HA-Cdt2^PIP-3A^-3FLAG. Proteasome inhibitor MG132 was used at 25 μM. UV-C (254 nm) irradiation of whole cells in dishes was performed at 5 to 100 J/m^2^ using a UV lamp (SUV-16, As One, Japan), a UV cross-linker (FS-800, Funakoshi) or a UV-box equipped with a UV-C lamp and a radiometer 254nm.

### Plasmids

Cdt2^WT(1-730)^-3FLAG expression plasmid, pCMV-HA-Cdt2^WT(1-730)^-3FLAG was constructed by cloning the PCR-amplified HA-Cdt2^WT(1-730)^ fragment into NcoI-NotI-cut pCMV-3FLAG. Cdt2^1-417^-3FLAG expression plasmid, pCMV-HA-Cdt2^1-417^-3FLAG and Cdt2^1-700^-3FLAG was constructed in a same way by PCR-amplifying the corresponding region. FLAG-Cdt2^300-730^, Cdt2^390-730^, and Cdt2^360-730^ expressing plasmids were constructed by PCR-amplifying the corresponding regions, (300-730, 390-730, and 600-730) and cloning into the p3FLAG-3NLS-myc plasmid which was prepared by cutting out the p21 gene from the p3FLAG-3NLS-myc-p21 plasmid (Nishitani et al., 2008). pcDNA-6myc-PCNA was constructed by ligating the PCR-amplified 6myc and PCNA. To construct the expression plasmid of PIP box mutant of Cdt2 (Cdt2^PIP-3A^-FLAG), pCMV-HA-Cdt2^PIP-3A^-3FLAG, Quick Change site-directed mutagenesis method (Strategene) was performed using primers: AGCTCCATGAGGAAAGCCTGCACAGCCGCCCATAGAAAGTCCCAGGAG and CTGGGACTTTCTATGGGCGGCTGTGCAGGCTTTCCTCATGGAGCTGGG. The baculovirus expression plasmids for HA-Cul4A and His-myc-Rbx1 were as described. The pBP8- Cdt2^WT(1-730)^-3FLAG plasmid was constructed by ligating the BamH1-Cdt2-Kpn1 fragment from pBP8-Cdt2 and the Kpn1-Cdt2-3FLAG-Sma1 fragment from pCMV-HA-Cdt2-3FLAG into pBacPAK8 between BamH1 and Sma1 sites. pBP9-DDB1 was constructed by ligating the PCR-amplified BamH1-DDB1-EcoR1 fragment and the EcoR1-DDB1-Xho1 fragment into pBacPAK9 at BamH1-Xho1 sites. pBP9-Cdt1-3FLAG was constructed by cloning the PCR amplified Sac1-Cdt1-3FLAG-Xho1 fragment using pCMV-Cdt1-3FLAG as a template into pBacPAK9. The GFP-Cdt2^wt^, GFP-Cdt2^(1-417)^ and GFP-Cdt2^(390-730)^ expression plasmids were constructed by subcloning of the respective FLAG-tagged constructs into pEGFP-C3 (Clontech). For the GFP-Cdt2^wt^ expression plasmid, pCMV-HA-Cdt2^WT(1-730)^-3FLAG was cut with NcoI and KpnI, KpnI and XmaI. In order to produce blunt ends for the NcoI sites, Klenow was used. The fragment was then cloned into the HindIII and XmaI sites of pEGFP-C3. The ends produced by HindIII were made blunt by Klenow. For the GFP-Cdt2^(1-417)^ the pCMV-HA-Cdt2^1-417^-3FLAG construct was cut by NcoI and XmaI. Klenow was used for the NcoI ends. The fragment was then cloned into the HindIII and XmaI sites of pEGFP-C3. The ends produced by HindIII were made blunt by Klenow. In order to achieve a better nuclear localization for the GFP-Cdt2^wt^ and GFP-Cdt2^(1-417)^ constructs these were sublconed into p3FLAG-3NLS-myc by XhoI and BamHI. Then they were subcloned again into pEGFP-C3 by using the HindIII-BamHI sites. For the GFP-Cdt2^(390-730)^ expression plasmid, the p3FLAG-3NLS-myc Cdt2^(390-730)^ construct was cut with HindIII and BamHI. The fragment produced was cloned into the HindIII-BamHI sites of pEGFP-C3.

### Antibodies, Western Blotting, and Immunofluorescence

For Western blotting, whole cell lysates were prepared by lysing cell pellets directly in SDS-PAGE buffer. For immunofluorescence, U2OS or HEK293 cells were fixed in 4% paraformaldehyde (WAKO) for 10 min, permeabilized in 0.25% (v/v) Triton X-100 in phosphate-buffered saline (PBS), and stained with the indicated antibodies as described previously. For double-staining with PCNA and Cdt2^WT^ or Cdt2^PIP-3A^, cells were permeabilized in 0.1% Triton X-100 for 1 min on ice, and were fixed in 4% paraformaldehyde (WAKO) for 10 min at room temperature, followed by fixation in ice-cold methanol for 10 min. For cyclobutane pyrimidine dimer (CPDs) staining, MCF7 and HeLa cells were fixed in 4% PFA for 10 minutes, permeabilized in 0.3% Triton X-100 in PBS and then incubated in 0.5 N NaOH for 5 min for DNA denaturation prior to immunofluorescence. Alexa 488-conjugated anti-mouse and Alexa592- conjugated anti-rabbit antibodies were used as secondary antibodies with Hoechst 33258 to visualize DNA. The following primary antibodies were used: Cdt1 (Nishitani et al., 2006), Cdt2 (Nishitani et al., 2008), cyclin A (mouse, Ab-6, Neomarkers; rabbit, H-432, Santa Cruz), Myc (mouse, 9E10, Santa Cruz), FLAG (F3165, F7425, SIGMA), PCNA (PC10, Santa Cruz; rabbit serum, a gift from Dr. Tsurimoto), XP-A (FL-273, Santa Cruz), RCC1(Shiomi et al., 2012a), DDB1(Bethyl Laboratories), Rbx1(ROC1, Ab-1, Neo Markers) and Cul4A (Bethyl Laboratories), DDB1 (Bethyl Laboratories), RFC1 (H-300, Santa Cruz Biotechnology), RFC4 (H-183, Santa Cruz Biotechnology) and DYKDDDDK Tag (Cell Signaling). CPDs(CosmoBio). Protein levels were analyzed by ImageJ software.

### Micropore UV-irradiation assay

We used a method to induce DNA photoproducts within localized areas of the cell nucleus as described in Ishii et al. To perform micropore UV-irradiation, cells were cultured on cover slips, washed twice with PBS and subsequently covered with an isopore polycarbonate membrane filter (Millipore) with a pore size of either 3 or 5 μm in diameter, and irradiated. UV-irradiation was achieved using a UV lamp (SUV-16, As One, Japan) with a dose rate of 0.4 J/m^2^•s, which was monitored with a UV radiometer (UVX Radiometer, UVP) at 254 μm, or with a UV box equipped with a UV-C lamp monitored with a VLX-3W radiometer (Vilber Lourmat) at 254 nm (CX-254 sensor). The filter was removed and cells were either fixed or cultured for the indicated time before fixation, and processed for immunofluorescence. For FRAP analysis, cells were cultured for 20 minutes to allow recruitment to sites of damage prior to photobleaching.

### Immunoprecipitation from Cdt2-FLAG expressing cells

Control or Cdt2-FLAG expressing 293 cells that were UV-irradiated (50 J/m^2^), not UV-irradiated, or in middle S phase (5 h after release from aphidicolon arrest) were fixed with 0.02% formaldehyde for 10 min, lysed using 0.1% Triton X-100 mCSK buffer (10 mM Pipes, pH 7.9, 100 mM NaCl, 300 mM sucrose, 0.1% (v/v) Triton X-100, 1 mM phenylmethylsulfonyl fluoride, 10 mM β-glycerophosphate, 1 mMNa_3_VO_4_, 10 mM NaF), and then sonicated. After centrifugation (30,000 rpm for 20 min at 4°C), the supernatants were mixed with anti-FLAG antibody-conjugated magnetic beads (M8823, SIGMA) for 1 hr at 4°C to obtain immunoprecipitates. The precipitate was washed with ice-cold 0.1% Triton X-100 containing mCSK buffer and subsequently suspended in SDS sample buffer.

### RNAi Knockdown Experiments

Double-stranded RNAs were transfected at 100 μM using Oligofectamine (Invitrogen) or HiPerFect (Qiagen). Twenty-four hours after the first transfection, a second transfection was performed and cells were cultured for two more days. The following siRNAs were made by Dharmacon: Skp2; GCAUGUACAGGUGGCUGUU, Cdt2_3’UTR; GCUGAGCUUUGGUCCACUA. The siRNA for siLuc, known as GL2, was used as a control siRNA. For Cdt1 silencing in Figure 1A and B, unsychronized HeLa cells were transfected twice with 200 nM of Cdt1 siRNA or control Luciferase siRNA using Lipofectamine 2000 with a time interval of 24 h and were analysed 48 hours after the second transfection. In synchronized MCF7 cells (Figure 1D) a single transfection with 200 nM of Cdt1 siRNA or control Luciferase siRNA using Lipofectamine 2000 5 hours after release from thymidine and before adding nocodazole was carried out. The Cdt1 siRNA was made by MWG: AACGUGGAUGAAGUACCCGACTT.

### Chromatin Fractionation

Cell extracts were prepared using 0.1% Triton X-100-containing mCSK buffer (200 µg of total protein in 400-800 µl). After centrifugation (15,000 rpm for 10 min at 4°C), soluble and chromatin-containing pellet fractions were obtained. The pellet was washed with ice-cold 0.1% Triton X-100 containing mCSK buffer and subsequently suspended in SDS sample buffer.

### Protein purification

*PCNA*: PCNA was purified essentially as a trimer as described (Fukuda et al., 1995). Cell lysate was prepared from pT7-PCNA transformed Rosetta(DE3) and sequentially subjected to Resource Q, Superdex 200, and Mono Q (GE Healthcare Life Science) column chromatography.

*Cdt1-3FLAG*: The baculoviruses for Cdt1-3FLAG expression were infected into Sf21 insect cells, and cultured at 27°C for 60 hours. Cell lysate was prepared with 0.5M NaCl containing Buffer B (50 mM Tris-HCl, pH8.0, 0.15 M NaCl, 1 mM EDTA, 10% glycerol, 1xProtease inhibitor cocktail (Roche Applied Science), 1 mM PMSF, 2 µg/mL Leupeptin), and passed on DEAE column. The flowthrough fraction was mixed with anti-FLAG resin (Sigma) and Cdt1-3FLAG was eluted with 200µg/mL 3xFLAG peptides (Sigma Aldrich).

*RFC complex*: The baculoviruses for FLAG-Rfc1, Rfc2, Rfc3, Rfc4 and Rfc5 were co-infected to High Five cells, expressed and purified as described(Shiomi et al., 2000). *CRL4^Cdt2^*; the baculoviruses for Cdt2-3FLAG (WT or PIP-3A), DDB1, HA-Cul4 and His-myc-Rbx1 were co-infected into Sf21 cells, and purifies as described (Hayashi et al., 2014). Briefly, the cell lysate was prepared with 0.5M NaCl and 0.5% NonidetP-40 containing Buffer B and clarified by centrifugation and passed on DEAE column. The flowthrough fraction was mixed with anti-FLAG resin and CRL4^Cdt2-3FLAG^ was eluted with Buffer B containing 0.1MNaCl, 0.1%NP-40 and 200 µg/mL 3XFLAG peptide (Sigma Aldrich). If necessary, CRL4^Cdt2-3FLAG^ was loaded on 15-35% glycerol gradient in buffer G (25 mM HEPES pH 7.8, 0.1 M NaCl, 1 mM EDTA, 0.01% NP-40) and subjected to ultra-centrifugation (Beckman TLS-55) at 214,000g for 18 hrs at 4 °C. 100 µl fractions were collected.

### *In vitro* binding assay for Cdt1 and CRL4^Cdt2^ with PCNA

Cdt1-3FLAG (WT) and CRL4^Cdt2-3FLAG (WT)^-beads were prepared as follows; (Figure 5B) Cleared lysates were made from insect cells infected with baculoviruses for Cdt1-3FLAG (WT) or CRL4^Cdt2-3FLAG (WT)^, incubated with anti-FLAG magnetic beads pre-blocked with Buffer B containing 50% FBS, 0.5 M NaCl and 0.5%NP-40 and washed with 0.5 M NaCl and 0.5%NP-40 containing Buffer B and then with 0.1 M NaCl and 0.1%NP-40 containing Buffer B. The protein beads prepared as above were incubated with PCNA protein at 4°C for two hours, washed and subjected to Western blotting.

### Preparation of nicked circular DNA-beads

Closed circular (cc) DNA-beads were prepared as described (Higashi et al., 2012). Briefly, single stranded DNA templates prepared using pBluescript II KS(-) was annealed with a biotinated oligonucleotide primer (5’-CGCCTTGATCGT [biotin-dT]GGGAACCGGAGCTGAATGAAGC-3’). After second strand synthesis and ligation, covalently closed circular double strand DNA was purified by CsCl/EtBr density gradient centrifugation, incubated with streptavidin and bound to biotin-sepharose beads to produce the cc DNA-beads (around 100ng per 1µl of bead). A single nick was introduced by Nb.BbvCI (New England Biolabs) treatment to produce the nicked circular (nc) DNA-beads. As control beads, biotin-sepharose beads were incubated with streptavidin alone (sa-beads).

### PCNA loading and binding assay with purified Cdt1 and CRL^Cdt2^

For loading of PCNA on DNA beads, 2 μl bed volume of nc DNA beads were incubated with 280 ng of RFC and 200 ng of PCNA in 20 μl of 2mM ATP containing HBS buffer (10 mM HEPES pH7.4, 10 mM MgCl_2_, 0.2 mM EDTA, 0.05 % Tween 20 and 0.15 M NaCl) at 37°C for one hour and washed with HBS buffer three times. The sa-beads were treated in a same way and used as control beads. Beads were incubated with Cdt1 or CRL4^Cdt2^ alone or with both Cdt1 and CRL4^Cdt2^ at 4 °C for one hour and washed with HBS buffer. The bound proteins were analyzed on Western blotting.

### *In vitro* binding of PIP box peptides to PCNA

The expression plasmid for full-length human PCNA has been described (Hibbert and Sixma, 2012). Human PCNA was expressed in Bl21 (DE3) *E. coli* cells (Merck KGaA, Darmstadt, Germany) and purified as described (Hibbert and Sixma, 2012). Peptides – Cdt1^PIP^ (704-717), Cdt1^PIP^-TAMRA (704-717), Cdt2^PIP^ (1-14) and Cdt2^PIP^-TAMRA (1-14) were synthesized in-house using Fmoc synthesis. Fluorescence polarization was monitored using a PheraStar plate reader (BMG Labtech) in non-binding 96 well plates (Corning), using an excitation wavelength of 540 nm and an emission wavelength of 590 nm. Experiments were performed in a buffer containing 10 mM HEPES pH 7.5, 125 mM NaCl and 0.5 mM Tris (2-carboxyethyl) phosphine hydrochloride. In the direct binding experiments, 1 nM of labelled peptide was mixed with PCNA at final concentrations ranging from 0 to 2 μM. Error bars represent standard deviations from triplicate experiments. The data were fitted according to standard equations (Littler et al., 2010).

### X-ray crystallography structure determination

The purified PCNA was co-crystallized with Cdt2 as well as Cdt1 peptides, using the sitting drop vapour diffusion method in MRC 3-Well Crystallization Plates (Swissci), with standard screening procedures (Newman et al., 2005). The protein solution at 17 mg mL^-1^ was pre-incubated with an equimolar amount of peptide, and 0.1 μL of this solution was mixed with 0.1 μL of reservoir solution and equilibrated against a 30 μL reservoir. Crystals for the Cdt2 complex were obtained in 10% (w/v) PEG 6000 and 100 mM MES/HCl, pH 6.0. Crystals for the Cdt1 complex were obtained in 15% (w/v) PEG 400, 100 mM calcium acetate and 100 mM MES/HCl, pH 6.0. Crystals appeared at 4 °C within 72 hours. Crystals were briefly transferred to a cryo-protectant solution containing the reservoir solution and 25% (w/v) PEG200 and vitrified by dipping in liquid nitrogen.

### Data collection and structure refinement

X-ray data were collected on beamline ID29 at the European Synchrotron Radiation Facility (ESRF). The images were integrated with XDS (Kabsch, 2010) and merged and scaled with AIMLESS (Evans, 2011). The starting phases were obtained by molecular replacement using PHASER (McCoy, 2007) with an available PCNA structure (PDB, 1AXC) as the search model. The models were built using COOT (Emsley et al., 2010) and refined with REFMAC (Murshudov et al., 2011) in iterative cycles. Model re-building and refinement parameter adjustment were performed in PDB-REDO (Joosten et al., 2014); homology-based hydrogen bond restraints (van Beusekom et al., 2018) were used at some stages of the procedure. The quality of the models was evaluated by MOLPROBITY (Chen et al., 2010). Data collection and refinement statistics are presented in Table 1.

### Fluorescence recovery after photobleaching (FRAP) experiments and data analysis

Cells (HeLa or MCF7) were plated on 35-mm glass-bottom dishes in phenol-red free medium (Invitrogen). FRAP experiments were conducted as previously described (Giakoumakis et al., 2017; Xouri et al., 2007b). A Leica TCS SP5 microscope equipped with a 63x magnification 1.4 NA oil-immersion lens and FRAP booster were used. During FRAP, cells were maintained at 37°C and 5% CO_2_. A defined circular region of interest with 2 μm diameter (ROI1) was placed in the nucleus and at the recruitment sites. GFP was excited using a 488 nm Argon laser line. 50 pre-bleach images were acquired with 3% of the 488 nm line at 60% Argon laser intensity, followed by a single bleach pulse on ROI1 using 476 nm and 488 nm laser lines combined at maximum power. In this manner at least 60% of the fluorescence in ROI1 was bleached successfully. Next, 400 post bleach images with a 0.052 sec interval were recorded. Mean intensities of ROI1, the whole nucleus (ROI2) and an area outside of the nucleus (ROI3) were quantified and exported as .csv format files. Data analysis was performed using easyFRAP (Giakoumakis et al., 2017; Koulouras et al., 2018; Rapsomaniki et al., 2012).

## Author contributions

Akiyo Hayashi:in vitro biochemical analysis, Cdt2-PIP analysis in cells and made figures, Nickolaos Nikiforos Giakoumakis: Cdt2 recruitment in cells, FRAP analysis, Tatjana Heidebrecht : produced recombinant proteins, performed in vitro biophysical assays and analysed data, and produced crystals, Takashi Ishii: Y2H screening and identification of PCNA and production of Cdt2 PIP-mutant and its initial analysis, Andreas Panagopoulos: Cdt2 N- and C- terminal recruitment in cells, Naohiro Suenaga: Cdt2-N-terminal and C-terminal domain analysis, Christophe Caillat: produced proteins, made initial experiments of PIP box binding, identified PIP box, Michiyo Takahara: isolation of U2OS stables and analysis, Richard G. Hibbert: identified the PIP box from sequence data, Tatsuro Takahashi: assistance for DNA-bead preparation, Magda Stadnik-Spiewak: technical assistance in protein production, Yasushi Shiomi: technical and analytical advice, Stavros Taraviras: supervised in cell analysis, Eleonore von Castelmur: collected and processed crystallographic data, supervised in vitro biophysics experiments, and solved PCNA complex structures, Zoi Lygerou: supervised recruitment and FRAP analysis in cells, contributed to ms writing, Anastassis Perrakis: prepared figures, refined crystal structures, wrote ms, supervised in vitro biophysics and structural work, Hideo Nishitani: supervised the biological characterization, wrote ms, and made figures

## Acknowledgements

This work was financially supported by JSPS KAKENHI Grant Numbers JP25131718 and JP26291025 (to HN), 25.7320 (to AH), the European Research Council (ERC-StG 281851 and ERC-PoC 755284), the Greek Secretariat for Research and Technology (Bioimaging-GR and Postdoctoral program) (to ZL) and a Greek state scholarship (to AP).

We thank the Advanced Light Microscopy Facility of the University of Patras.

## Supplementary Figure legend

**Figure S1 A. FRAP analysis of Cdt2 binding to sites of damage in the absence of Cdt1**

Quantitative parameters, Immobile fraction and half-maximal recovery time of the mobile fraction (T^1/2^), corresponding to the FRAP curves described in figure 1C and showing the recovery of Cdt1-eGFP, Cdt2-eGFP, PCNA-eGFP and eGFP-NSL as a control, in undamaged and locally UV-C damaged cells. Mean values with standard deviation, as well as the number of individual cells analysed by FRAP in each case are shown. Normalization and Curve fitting of individual FRAP curves for parameter estimation was performed with easyFRAP. **B.** FRAP images before, during and after photobleaching at the sites of damage. **C.** Mean normalized recovery curves of Cdt1-eGFP, Cdt2-eGFP, PCNA-eGFP and eGFP-NSL in undamaged cells. **D.** Efficiency of synchronization and Cdt1 depletion of MCF7 cells. The percentage of cells expressing Cdt1 or cyclin A is shown for MCF7 cells synchronized in G1, without (siLuc) or with (siCdt1) Cdt1 depletion, as described in figure 1D. **E.** Quantitative parameters, Immobile fraction and half-maximal recovery time of the mobile fraction (T^1/2^), corresponding to the FRAP curves described in figure 1D and showing Cdt2 in the presence and absence of Cdt1 in undamaged and locally UV-C damaged cells. Mean values with standard deviation, as well as the number of individual cells analysed by FRAP in each case are shown. Normalization and Curve fitting of individual FRAP curves was done with easyFRAP.

**Figure S2.** Yeast two hybrid screening identified PCNA as a Cdt2 C-terminus interacting protein.

**Figure S3.** crystallography: Electron density maps showing the Cdt1 and Cdt1 peptides electron density after structure refinement. The 2mF_o_-DF_c_ map is contoured to 1.0 rms only around the modelled peptides. A characteristic monomer from the PCNA trimer is show is each case.

**Figure S4. Purified proteins used for *in vitro* assay.**

**A**. Purified Cdt1-3FLAG, PCNA and RFC proteins. **B**. CRL4^Cdt2^ complex after purification on FLAG-resins. The asterisks are contaminants. The purified proteins were Western blotted with antibodies for each protein. **C**. The CRL4^Cdt2^ complex after purification on FLAG-resins was further purified on glycerol gradient centrifugation. The peak fractions of CRL4^Cdt2^ tetramer (calculated molecular weight, 313 kDa) were indicated by the dotted line.

**Figure S5. Loading of PCNA on nicked circular DNA-beads and following binding assay.**

**A.** Scheme for PCNA loading on nicked circular (nc) DNA and for following Cdt1 and CRL4^Cdt2^ binding assay. Nicked circular DNA beads or control beads were incubated with PCNA and RFC for 1 hr at 37 °C and were washed (Step1; Loading of PCNA). The beads were then incubated with Cdt1 or CRL4^Cdt2^ (Step2; Binding). **B**. RFC dependent loading of PCNA.

**Figure S6. Cdt2 PIP-box is important for its recruiting to DNA replication sites and its poly-ubiquitination activity in HEK293 cells.**

A. HEK293 cells or cells stable Cdt2WT or Cdt2PIP-3A were pre-extracted, fixed and stained with PCNA and DYKDDDDK (FLAG) antibodies. **B**. Poly-ubiquitination after UV-irradiation. HEK293 and its stables expressing Cdt2^WT^ or Cdt2^PIP-3A^ were transfected with siCdt2 or control (siLuc). Three days later, cells were treated with MG132 (25 μM), and 10 min later irradiated with UV (20 J/m^2^). Cells were collected at the indicated time points, and used for Western blotting with indicated antibodies.

**Figure S7. Cdt2 PIP-box is important for CRL4^Cdt2^ activity in U2OS cells.**

**A**. U2OS stable cells expressing Cdt2^WT^ or Cdt2^PIP-3A^. U2OS cells were transfected with expression vectors for Cdt2^WT^-FLAG or Cdt2^PIP-3A^-FLAG, and isolated stable clones were verified by immunofluorescent analysis and Western blot analysis. Frequency of expressing cells for each construct was shown (%). **B**. Poly-ubiquitination assay with U2OS stable cells. U2OS cells and Cdt2^WT^-3FLAG or Cdt2^PIP-3A^-3FLAG expressing stable U2OS cells were treated with MG132 for 3 hr or not (-), and whole cell extracts were prepared for Western blotting. **C.** Cells were transfected with siCdt2 or control siLuc for 72 hr, irradiated with UV (+, 5 J/m^2^) or not (-) and collected 1hr later for Western blotting. Relative levels of Cdt1 were shown, set the UV (-) level as 1.0 (average of two independent experiments).

**Figure S8. Cdt2^PIP-3A^ mutant shows enlarged nuclei.**

Cells transfected with siLuc or siCdt2 for 72 hr were stained with Hoechst 33258.

## REFERENCES

Abbas, T., and Dutta, A. (2011). CRL4Cdt2: master coordinator of cell cycle progression and genome stability. Cell cycle (Georgetown, Tex) 10, 241-249.

Abbas, T., Mueller, A.C., Shibata, E., Keaton, M., Rossi, M., and Dutta, A. (2013). CRL1-FBXO11 Promotes Cdt2 Ubiquitylation and Degradation and Regulates Pr-Set7/Set8-Mediated Cellular Migration. Molecular cell.

Abbas, T., Sivaprasad, U., Terai, K., Amador, V., Pagano, M., and Dutta, A. (2008). PCNA-dependent regulation of p21 ubiquitylation and degradation via the CRL4Cdt2 ubiquitin ligase complex. Genes & development 22, 2496-2506.

Arias, E.E., and Walter, J.C. (2006). PCNA functions as a molecular platform to trigger Cdt1 destruction and prevent re-replication. Nat Cell Biol 8, 84-90.

Arias, E.E., and Walter, J.C. (2007). Strength in numbers: preventing rereplication via multiple mechanisms in eukaryotic cells. Genes & development 21, 497-518.

Bell, S.P., and Dutta, A. (2002). DNA replication in eukaryotic cells. Annual review of biochemistry 71, 333-374.

Blow, J.J., and Dutta, A. (2005). Preventing re-replication of chromosomal DNA. Nature reviews Molecular cell biology 6, 476-486.

Branzei, D., and Foiani, M. (2010). Maintaining genome stability at the replication fork. Nature reviews Molecular cell biology 11, 208-219.

Bruning, J.B., and Shamoo, Y. (2004). Structural and thermodynamic analysis of human PCNA with peptides derived from DNA polymerase-delta p66 subunit and flap endonuclease-1. Structure 12, 2209-2219.

Cardozo, T., and Pagano, M. (2004). The SCF ubiquitin ligase: insights into a molecular machine. Nature reviews Molecular cell biology 5, 739-751.

Centore, R.C., Havens, C.G., Manning, A.L., Li, J.M., Flynn, R.L., Tse, A., Jin, J., Dyson, N.J., Walter, J.C., and Zou, L. (2010). CRL4(Cdt2)-mediated destruction of the histone methyltransferase Set8 prevents premature chromatin compaction in S phase. Molecular cell 40, 22-33.

Chen, V.B., Arendall, W.B., 3rd, Headd, J.J., Keedy, D.A., Immormino, R.M., Kapral, G.J., Murray, L.W., Richardson, J.S., and Richardson, D.C. (2010). MolProbity: all-atom structure validation for macromolecular crystallography. Acta Crystallogr D Biol Crystallogr 66, 12-21.

Clijsters, L., and Wolthuis, R. (2014). PIP-box-mediated degradation prohibits re-accumulation of Cdc6 during S phase. Journal of cell science 127, 1336-1345.

De Biasio, A., Campos-Olivas, R., Sanchez, R., Lopez-Alonso, J.P., Pantoja-Uceda, D., Merino, N., Villate, M., Martin-Garcia, J.M., Castillo, F., Luque, I., et al. (2012). Proliferating cell nuclear antigen (PCNA) interactions in solution studied by NMR. PLoS One 7, e48390.

Diffley, J.F. (2004). Regulation of early events in chromosome replication. Current biology : CB 14, R778-786.

Emsley, P., Lohkamp, B., Scott, W.G., and Cowtan, K. (2010). Features and development of Coot. Acta Crystallogr D Biol Crystallogr 66, 486-501.

Evans, P.R. (2011). An introduction to data reduction: space-group determination, scaling and intensity statistics. Acta Crystallogr D Biol Crystallogr 67, 282-292.

Fischer, E.S., Scrima, A., Bohm, K., Matsumoto, S., Lingaraju, G.M., Faty, M., Yasuda, T., Cavadini, S., Wakasugi, M., Hanaoka, F., et al. (2011). The molecular basis of CRL4DDB2/CSA ubiquitin ligase architecture, targeting, and activation. Cell 147, 1024-1039.

Fukuda, K., Morioka, H., Imajou, S., Ikeda, S., Ohtsuka, E., and Tsurimoto, T. (1995). Structure-function relationship of the eukaryotic DNA replication factor, proliferating cell nuclear antigen. The Journal of biological chemistry 270, 22527-22534.

Giakoumakis, N.N., Rapsomaniki, M.A., and Lygerou, Z. (2017). Analysis of Protein Kinetics Using Fluorescence Recovery After Photobleaching (FRAP). Methods Mol Biol 1563, 243-267.

Han, C., Wani, G., Zhao, R., Qian, J., Sharma, N., He, J., Zhu, Q., Wang, Q.E., and Wani, A.A. (2015). Cdt2-mediated XPG degradation promotes gap-filling DNA synthesis in nucleotide excision repair. Cell cycle (Georgetown, Tex) 14, 1103-1115.

Havens, C.G., Shobnam, N., Guarino, E., Centore, R.C., Zou, L., Kearsey, S.E., and Walter, J.C. (2012). Direct Role for proliferating cell nuclear antigen (PCNA) in substrate recognition by the E3 Ubiquitin ligase CRL4-Cdt2. The Journal of biological chemistry.

Havens, C.G., and Walter, J.C. (2009). Docking of a specialized PIP Box onto chromatin-bound PCNA creates a degron for the ubiquitin ligase CRL4Cdt2. Molecular cell 35, 93-104.

Havens, C.G., and Walter, J.C. (2011). Mechanism of CRL4Cdt2, a PCNA-dependent E3 ubiquitin ligase. Genes & development 25, 1568-1582.

Hayashi, A., Suenaga, N., Shiomi, Y., and Nishitani, H. (2014). PCNA-dependent ubiquitination of Cdt1 and p21 in mammalian cells. Methods Mol Biol 1170, 367-382.

He, Y.J., McCall, C.M., Hu, J., Zeng, Y., and Xiong, Y. (2006). DDB1 functions as a linker to recruit receptor WD40 proteins to CUL4-ROC1 ubiquitin ligases. Genes & development 20, 2949-2954.

Hibbert, R.G., and Sixma, T.K. (2012). Intrinsic flexibility of ubiquitin on proliferating cell nuclear antigen (PCNA) in translesion synthesis. The Journal of biological chemistry 287, 39216-39223.

Higa, L.A., Banks, D., Wu, M., Kobayashi, R., Sun, H., and Zhang, H. (2006). L2DTL/CDT2 interacts with the CUL4/DDB1 complex and PCNA and regulates CDT1 proteolysis in response to DNA damage. Cell cycle (Georgetown, Tex) 5, 1675-1680.

Higashi, T.L., Ikeda, M., Tanaka, H., Nakagawa, T., Bando, M., Shirahige, K., Kubota, Y., Takisawa, H., Masukata, H., and Takahashi, T.S. (2012). The prereplication complex recruits XEco2 to chromatin to promote cohesin acetylation in Xenopus egg extracts. Current biology : CB 22, 977-988.

Hoeijmakers, J.H. (2001). Genome maintenance mechanisms for preventing cancer. Nature 411, 366-374.

Hofmann, J.F., and Beach, D. (1994). cdt1 is an essential target of the Cdc10/Sct1 transcription factor: requirement for DNA replication and inhibition of mitosis. Embo J 13, 425-434.

Ishii, T., Shiomi, Y., Takami, T., Murakami, Y., Ohnishi, N., and Nishitani, H. (2010). Proliferating cell nuclear antigen-dependent rapid recruitment of Cdt1 and CRL4Cdt2 at DNA-damaged sites after UV irradiation in HeLa cells. The Journal of biological chemistry 285, 41993-42000.

Ivanov, I., Chapados, B.R., McCammon, J.A., and Tainer, J.A. (2006). Proliferating cell nuclear antigen loaded onto double-stranded DNA: dynamics, minor groove interactions and functional implications. Nucleic acids research 34, 6023-6033.

Jackson, S., and Xiong, Y. (2009). CRL4s: the CUL4-RING E3 ubiquitin ligases. Trends Biochem Sci 34, 562-570.

Jin, J., Arias, E.E., Chen, J., Harper, J.W., and Walter, J.C. (2006). A family of diverse Cul4-Ddb1-interacting proteins includes Cdt2, which is required for S phase destruction of the replication factor Cdt1. Molecular cell 23, 709-721.

Joosten, R.P., Long, F., Murshudov, G.N., and Perrakis, A. (2014). The PDB_REDO server for macromolecular structure model optimization. IUCrJ 1, 213-220.

Jorgensen, S., Eskildsen, M., Fugger, K., Hansen, L., Larsen, M.S., Kousholt, A.N., Syljuasen, R.G., Trelle, M.B., Jensen, O.N., Helin, K., et al. (2011). SET8 is degraded via PCNA-coupled CRL4(CDT2) ubiquitylation in S phase and after UV irradiation. J Cell Biol 192, 43-54.

Kabsch, W. (2010). Xds. Acta Crystallogr D Biol Crystallogr 66, 125-132.

Kim, D.H., Budhavarapu, V.N., Herrera, C.R., Nam, H.W., Kim, Y.S., and Yew, P.R. (2010). The CRL4Cdt2 ubiquitin ligase mediates the proteolysis of cyclin-dependent kinase inhibitor Xic1 through a direct association with PCNA. Molecular and cellular biology 30, 4120-4133.

Kim, Y., and Kipreos, E.T. (2007). The Caenorhabditis elegans replication licensing factor CDT-1 is targeted for degradation by the CUL-4/DDB-1 complex. Molecular and cellular biology 27, 1394-1406.

Kim, Y., Starostina, N.G., and Kipreos, E.T. (2008). The CRL4Cdt2 ubiquitin ligase targets the degradation of p21Cip1 to control replication licensing. Genes & development 22, 2507-2519.

Koulouras, G., Panagopoulos, A., Rapsomaniki, M.A., Giakoumakis, N.N., Taraviras, S., and Lygerou, Z. (2018). EasyFRAP-web: a web-based tool for the analysis of fluorescence recovery after photobleaching data. Nucleic acids research 46, W467-W472.

Li, X., Zhao, Q., Liao, R., Sun, P., and Wu, X. (2003). The SCF(Skp2) ubiquitin ligase complex interacts with the human replication licensing factor Cdt1 and regulates Cdt1 degradation. The Journal of biological chemistry 278, 30854-30858.

Littler, D.R., Alvarez-Fernandez, M., Stein, A., Hibbert, R.G., Heidebrecht, T., Aloy, P., Medema, R.H., and Perrakis, A. (2010). Structure of the FoxM1 DNA-recognition domain bound to a promoter sequence. Nucleic acids research 38, 4527-4538.

Mansilla, S.F., Soria, G., Vallerga, M.B., Habif, M., Martinez-Lopez, W., Prives, C., and Gottifredi, V. (2013). UV-triggered p21 degradation facilitates damaged-DNA replication and preserves genomic stability. Nucleic acids research 41, 6942-6951.

McCoy, A.J., Grosse-Kunstleve, R.W., Adams, P.D., Winn, M.D., Storoni, L.C., and Read, R.J. (2007). Phaser crystallographic software. Journal of applied crystallography 40, 658-674.

Michishita, M., Morimoto, A., Ishii, T., Komori, H., Shiomi, Y., Higuchi, Y., and Nishitani, H. (2011). Positively charged residues located downstream of PIP box, together with TD amino acids within PIP box, are important for CRL4(Cdt2)-mediated proteolysis. Genes to cells : devoted to molecular & cellular mechanisms 16, 12-22.

Murshudov, G.N., Skubak, P., Lebedev, A.A., Pannu, N.S., Steiner, R.A., Nicholls, R.A., Winn, M.D., Long, F., and Vagin, A.A. (2011). REFMAC5 for the refinement of macromolecular crystal structures. Acta Crystallogr D Biol Crystallogr 67, 355-367.

Newman, J., Egan, D., Walter, T.S., Meged, R., Berry, I., Ben Jelloul, M., Sussman, J.L., Stuart, D.I., and Perrakis, A. (2005). Towards rationalization of crystallization screening for small- to medium-sized academic laboratories: the PACT/JCSG+ strategy. Acta Crystallogr D Biol Crystallogr 61, 1426-1431.

Nishitani, H., and Lygerou, Z. (2004). DNA replication licensing. Front Biosci 9, 2115-2132.

Nishitani, H., Shiomi, Y., Iida, H., Michishita, M., Takami, T., and Tsurimoto, T. (2008). CDK inhibitor p21 is degraded by a proliferating cell nuclear antigen-coupled Cul4-DDB1Cdt2 pathway during S phase and after UV irradiation. The Journal of biological chemistry 283, 29045-29052.

Nishitani, H., Sugimoto, N., Roukos, V., Nakanishi, Y., Saijo, M., Obuse, C., Tsurimoto, T., Nakayama, K.I., Nakayama, K., Fujita, M., et al. (2006). Two E3 ubiquitin ligases, SCF-Skp2 and DDB1-Cul4, target human Cdt1 for proteolysis. EMBO J 25, 1126-1136.

Nukina, K., Hayashi, A., Shiomi, Y., Sugasawa, K., Ohtsubo, M., and Nishitani, H. (2018). Mutations at multiple CDK phosphorylation consensus sites on Cdt2 increase the affinity of CRL4(Cdt2) for PCNA and its ubiquitination activity in S phase. Genes to cells : devoted to molecular & cellular mechanisms 23, 200-213.

Oda, H., Hubner, M.R., Beck, D.B., Vermeulen, M., Hurwitz, J., Spector, D.L., and Reinberg, D. (2010). Regulation of the histone H4 monomethylase PR-Set7 by CRL4(Cdt2)-mediated PCNA-dependent degradation during DNA damage. Molecular cell 40, 364-376.

Petroski, M.D., and Deshaies, R.J. (2005). Function and regulation of cullin-RING ubiquitin ligases. Nature reviews Molecular cell biology 6, 9-20.

Ralph, E., Boye, E., and Kearsey, S.E. (2006). DNA damage induces Cdt1 proteolysis in fission yeast through a pathway dependent on Cdt2 and Ddb1. EMBO reports 7, 1134-1139.

Raman, M., Havens, C.G., Walter, J.C., and Harper, J.W. (2011). A genome-wide screen identifies p97 as an essential regulator of DNA damage-dependent CDT1 destruction. Molecular cell 44, 72-84.

Rapsomaniki, M.A., Cinquemani, E., Giakoumakis, N.N., Kotsantis, P., Lygeros, J., and Lygerou, Z. (2015). Inference of protein kinetics by stochastic modeling and simulation of fluorescence recovery after photobleaching experiments. Bioinformatics 31, 355-362.

Rapsomaniki, M.A., Kotsantis, P., Symeonidou, I.E., Giakoumakis, N.N., Taraviras, S., and Lygerou, Z. (2012). easyFRAP: an interactive, easy-to-use tool for qualitative and quantitative analysis of FRAP data. Bioinformatics 28, 1800-1801.

Rizzardi, L.F., Coleman, K.E., Varma, D., Matson, J.P., Oh, S., and Cook, J.G. (2015). CDK1-dependent inhibition of the E3 ubiquitin ligase CRL4CDT2 ensures robust transition from S Phase to Mitosis. The Journal of biological chemistry 290, 556-567.

Roukos, V., Kinkhabwala, A., Colombelli, J., Kotsantis, P., Taraviras, S., Nishitani, H., Stelzer, E., Bastiaens, P., and Lygerou, Z. (2011). Dynamic recruitment of licensing factor Cdt1 to sites of DNA damage. Journal of cell science 124, 422-434.

Sakaguchi, H., Takami, T., Yasutani, Y., Maeda, T., Morino, M., Ishii, T., Shiomi, Y., and Nishitani, H. (2012). Checkpoint kinase ATR phosphorylates Cdt2, a substrate receptor of CRL4 ubiquitin ligase, and promotes the degradation of Cdt1 following UV irradiation. PLoS One 7, e46480.

Sansam, C.L., Shepard, J.L., Lai, K., Ianari, A., Danielian, P.S., Amsterdam, A., Hopkins, N., and Lees, J.A. (2006). DTL/CDT2 is essential for both CDT1 regulation and the early G2/M checkpoint. Genes & development 20, 3117-3129.

Scrima, A., Konickova, R., Czyzewski, B.K., Kawasaki, Y., Jeffrey, P.D., Groisman, R., Nakatani, Y., Iwai, S., Pavletich, N.P., and Thoma, N.H. (2008). Structural basis of UV DNA-damage recognition by the DDB1-DDB2 complex. Cell 135, 1213-1223.

Senga, T., Sivaprasad, U., Zhu, W., Park, J.H., Arias, E.E., Walter, J.C., and Dutta, A. (2006). PCNA is a cofactor for Cdt1 degradation by CUL4/DDB1-mediated N-terminal ubiquitination. The Journal of biological chemistry 281, 6246-6252.

Shibata, E., Dar, A., and Dutta, A. (2014). CRL4Cdt2 E3 ubiquitin ligase and proliferating cell nuclear antigen (PCNA) cooperate to degrade thymine DNA glycosylase in S phase. The Journal of biological chemistry 289, 23056-23064.

Shiomi, Y., Hayashi, A., Ishii, T., Shinmyozu, K., Nakayama, J., Sugasawa, K., and Nishitani, H. (2012a). Two different replication factor C proteins, Ctf18 and RFC1, separately control PCNA-CRL4Cdt2-mediated Cdt1 proteolysis during S phase and following UV irradiation. Molecular and cellular biology 32, 2279-2288.

Shiomi, Y., Hayashi, A., Ishii, T., Shinmyozu, K., Nakayama, J.I., Sugasawa, K., and Nishitani, H. (2012b). Two different RFC proteins, Ctf18 and RFC1 separately control PCNA-CRL4Cdt2-mediated Cdt1 proteolysis during S phase and following UV-irradiation. Mol Cell Biol.

Shiomi, Y., Usukura, J., Masamura, Y., Takeyasu, K., Nakayama, Y., Obuse, C., Yoshikawa, H., and Tsurimoto, T. (2000). ATP-dependent structural change of the eukaryotic clamp-loader protein, replication factor C. Proceedings of the National Academy of Sciences of the United States of America 97, 14127-14132.

Slenn, T.J., Morris, B., Havens, C.G., Freeman, R.M., Jr., Takahashi, T.S., and Walter, J.C. (2014). Thymine DNA glycosylase is a CRL4Cdt2 substrate. The Journal of biological chemistry 289, 23043-23055.

Stathopoulou, A., Roukos, V., Petropoulou, C., Kotsantis, P., Karantzelis, N., Nishitani, H., Lygerou, Z., and Taraviras, S. (2012). Cdt1 is differentially targeted for degradation by anticancer chemotherapeutic drugs. PLoS One 7, e34621.

Sugimoto, N., Tatsumi, Y., Tsurumi, T., Matsukage, A., Kiyono, T., Nishitani, H., and Fujita, M. (2004). Cdt1 phosphorylation by cyclin A-dependent kinases negatively regulates its function without affecting geminin binding. The Journal of biological chemistry 279, 19691-19697.

Symeonidou, I.E., Taraviras, S., and Lygerou, Z. (2012). Control over DNA replication in time and space. FEBS letters 586, 2803-2812.

Tanaka, M., Takahara, M., Nukina, K., Hayashi, A., Sakai, W., Sugasawa, K., Shiomi, Y., and Nishitani, H. (2017). Mismatch repair proteins recruited to ultraviolet light-damaged sites lead to degradation of licensing factor Cdt1 in the G1 phase. Cell cycle (Georgetown, Tex) 16, 673-684.

Tardat, M., Brustel, J., Kirsh, O., Lefevbre, C., Callanan, M., Sardet, C., and Julien, E. (2010). The histone H4 Lys 20 methyltransferase PR-Set7 regulates replication origins in mammalian cells. Nat Cell Biol 12, 1086-1093.

Terai, K., Shibata, E., Abbas, T., and Dutta, A. (2013). Degradation of p12 subunit by CRL4Cdt2 E3 ligase inhibits fork progression after DNA damage. The Journal of biological chemistry 288, 30509-30514.

Tsakraklides, V., and Bell, S.P. (2010). Dynamics of pre-replicative complex assembly. The Journal of biological chemistry 285, 9437-9443.

Tsanov, N., Kermi, C., Coulombe, P., Van der Laan, S., Hodroj, D., and Maiorano, D. (2014). PIP degron proteins, substrates of CRL4Cdt2, and not PIP boxes, interfere with DNA polymerase eta and kappa focus formation on UV damage. Nucleic acids research 42, 3692-3706.

van Beusekom, B., Touw, W.G., Tatineni, M., Somani, S., Rajagopal, G., Luo, J., Gilliland, G.L., Perrakis, A., and Joosten, R.P. (2018). Homology-based hydrogen bond information improves crystallographic structures in the PDB. Protein Sci 27, 798-808. crystallographica Section D, Biological crystallography 67, 235-242.

Xouri, G., Dimaki, M., Bastiaens, P.I., and Lygerou, Z. (2007a). Cdt1 interactions in the licensing process: a model for dynamic spatiotemporal control of licensing. Cell cycle (Georgetown, Tex) 6, 1549-1552.

Xouri, G., Squire, A., Dimaki, M., Geverts, B., Verveer, P.J., Taraviras, S., Nishitani, H., Houtsmuller, A.B., Bastiaens, P.I., and Lygerou, Z. (2007b). Cdt1 associates dynamically with chromatin throughout G1 and recruits Geminin onto chromatin. EMBO J 26, 1303-1314.

